# Beyond brain size

**DOI:** 10.1101/145334

**Authors:** Corina J Logan, Shahar Avin, Neeltje Boogert, Andrew Buskell, Fiona R. Cross, Adrian Currie, Sarah Jelbert, Dieter Lukas, Rafael Mares, Ana F Navarrete, Shuichi Shigeno, Stephen H Montgomery

## Abstract

Despite prolonged interest in comparing brain size and behavioral proxies of ‘intelligence’ across taxa, the adaptive and cognitive significance of brain size variation remains elusive. Central to this problem is the continued focus on hominid cognition as a benchmark, and the assumption that behavioral complexity has a simple relationship with brain size. Although comparative studies of brain size have been criticized for not reflecting how evolution actually operates, and for producing spurious, inconsistent results, the causes of these limitations have received little discussion. We show how these issues arise from implicit assumptions about what brain size measures and how it correlates with behavioral and cognitive traits. We explore how inconsistencies can arise through heterogeneity in evolutionary trajectories and selection pressures on neuroanatomy or neurophysiology across taxa. We examine how interference from ecological and life history variables complicates interpretations of brain-behavior correlations, and point out how this problem is exacerbated by the limitations of brain and cognitive measures. These considerations, and the diversity of brain morphologies and behavioral capacities, suggest that comparative brain-behavior research can make greater progress by focusing on specific neuroanatomical and behavioral traits within relevant ecological and evolutionary contexts. We suggest that a synergistic combination of the ‘bottom up’ approach of classical neuroethology and the ‘top down’ approach of comparative biology/psychology within closely related but behaviorally diverse clades can limit the effects of heterogeneity, interference, and noise. We argue this shift away from broad-scale analyses of superficial phenotypes will provide deeper, more robust insights into brain evolution.

## 1) Motivation

In 1856, unusual bones were unearthed in a German limestone quarry in the Neander valley. Though later identified as the first Neanderthal specimen, the initial identification of the bones as a distinct species was highly contested (Madison, 2016). Indeed, the unusual size and features of the skull led to a range of questions: might the bones be from a pathological human or a new variety of fossil ape? One of the key novel methods employed to answer these questions was ‘craniometry’: the measurement of morphological features of the skull and their relationships (Goodrum, 2016; Madison, 2016). Employing these methods, Schaaffhausen (1858) found that the skull likely housed a large brain, falling within the range of contemporary humans. To many scientists at the time (though not all), the fossil’s brain size was evidence enough for the human, or human-like, status of the fossils (cf. Madison, 2016).

The distinctive brain morphology—and associated behavioral features—of hominins, and *Homo sapiens* in particular, continue to fascinate. Indeed, comparisons between human and non-human brains remain a central investigative target in contemporary comparative research. Such investigations have expanded the brain measurement tool-kit: in addition to craniometry, metrics such as brain size relative to body size (e.g., encephalization quotients), absolute brain size, and neuronal density are now common (Montgomery, 2017).

One of the central motivations for the continued research into brain measurement is its potential to reveal links between neuroanatomical structures and cognitive capabilities. Yet, just as debates about the special status of Neanderthals hinged upon the size of its cranium relative to humans, contemporary debates on the evolution of brain size and complex behavior have tended to privilege measures where humans come out on top. This bias has been built into a number of ‘monolithic’ general hypotheses (Barton, 2012) that claim links between measures of brain size and a diverse range of proxy-measures of complex behavior, such as ‘social’ intelligence (Dunbar & Shultz, 2007a, 2007b), ‘cultural’ intelligence (Tomasello, 1999; van Schaik & Burkart, 2011; van Schaik, Isler, & Burkart, 2012), ‘general’ intelligence (Burkart, Schubiger, & van Schaik, 2016; Reader, Hager, & Laland, 2011), and behavioral drive (Navarrete, Reader, Street, Whalen, & Laland, 2016; Wyles, Kunkel, & Wilson, 1983). In each of these cases, *Homo sapiens* emerges as the presumed pinnacle of a trajectory of brain evolution that correlates with increasing behavioral flexibility, intelligence, or socialization. Yet, both the significance of brain size and the interpretation of the correlated behaviors as more ‘complex’ or ‘cognitive’ remain poorly elucidated.

Here, we build on arguments made by Healy and Rowe (2007), unpack problems associated with using proxies, bring recent evidence from molecular techniques into the debate, and develop a framework that incorporates bottom-up and top-down approaches to advance the field. We argue that a more fruitful approach to linking brain measures and cognition is to de-emphasize coarse-grained notions of ‘intelligence’ and whole-brain measurements in favor of i) taxa-specific measurements of brains and ecologically meaningful behaviors, and ii) ‘bottom-up’ extrapolation of intraspecies measures based on phylogenetic context. This means measuring ecologically relevant features of brains and behaviors directly, rather than using proxies, at the within-species level. Then comparing closely related species to understand the relationships between traits, which will inform the even broader taxonomic scaling to make cross-species generalizations based on these validated correlations. Central to this is a movement away from *Homo sapiens* as the measuring stick for evaluating the neuroanatomical features and behavioral capabilities of other animals.

Below, we introduce a wide variety of research that examines brains and behavior across various phyla and discuss how lessons learned from disparate taxa can inform the way we interpret brain evolution, even among more familiar taxa such as vertebrates. Our aim is to emphasize the advantages and disadvantages of the different metrics, methods, and assumptions in this field. We review criticisms leveled against comparative studies of brain size, but go further by establishing why the recognized limitations arise. By doing so, we show why broad-stroke narratives struggle to capture the wide diversity of neuroanatomical features and behavioral capacities in animals. As a result, we argue that a more targeted ‘bottom-up’ approach that measures brains and behaviors at the intraspecies level to investigate cognitive, neuroanatomical, and behavioral diversity is needed to fully understand how behavioral complexity emerges from neural systems, and how well, or poorly, brain size reflects this variation.

## 2) Limitations of research on brain size and cognition

Interpreting how variation in brain size might be related to variation in cognition involves a set of assumptions that are frequently made in comparative studies:

- Brain size can be measured with negligible error
- Investing in a larger brain comes at a cost of investing in other tissues and/or life history traits
- Scaling relationships between brain size and body size are conserved within and across species
- Brain regions scale uniformly with total brain size
- Brain size scales with neuron number
- Cognitive abilities are discretely coded in the brain
- Cognitive abilities can be unambiguously ascertained by measuring behavior
- Brain size is directly and linearly associated with variation in cognition
- Selection on cognitive abilities and brain measures acts uniformly across species

These assumptions are applied uniformly both across and within species. The validity of these assumptions has previously been challenged by Snell (1892) and Healy and Rowe (2007), and we provide additional arguments in this section. First, the use of brain size as a trait makes implicit assumptions about how brains develop and evolve (see §2.1). Second, when correlating brain size and a measure of cognition we make assumptions about how selection acts on, or for, either trait (see §2.2). Finally, measuring cognition inevitably requires making some assumptions about the nature of behavioral complexity and what we view as a ‘cognitive’ trait (see §2.3). In each case, the lack of data supporting the validity of these assumptions directly limits our capacity to make reliable inferences about the link between brain size and cognition.

### 2.1 Assumptions and limitations of what brain size measures

Brain size may seem an easy neuroanatomical trait to measure, and the ease of obtaining a data point for a species, using one to a few specimens, renders it a historically useful starting point for many studies (Healy & Rowe, 2007; Jerison, 1985). However, brain size has also become the end point for many studies, with the variability of this trait becoming a target for evolutionary explanation. Large databases are populated by both individual measures and species’ brain size averages, which are used to examine cross-species correlations between brain size and a number of other traits. Researchers look to these databases for answers to questions such as *What is the significance of a large brain? What are the costs, and what are the benefits?* (e.g., Aiello & Wheeler, 1995; Armstrong, 1983; Harvey & Bennett, 1983; Isler & van Schaik, 2009; Nyberg, 1971). Cross-species correlations reveal that relative brain size (brain size relative to body size) is putatively associated with a range of life history and ecological traits. For example, relative brain size may correlate positively with longevity (a benefit) and negatively with fecundity (a cost) in mammals (Allman, McLaughlin, & Hakeem, 1993; Deaner, Barton, & van Schaik, 2003; González-Lagos, Sol, & Reader, 2010; Isler, 2011; Isler & van Schaik, 2009; Sol, Székely, Liker, & Lefebvre, 2007). Crucially, however, these correlations are not necessarily independent nor consistent across taxa; for example, relative brain size and longevity do not significantly correlate in strepsirrhine primates (lemurs and lorises; Allman et al., 1993). Other analyses suggest the relationship may be a consequence of developmental costs rather than an adaptive relationship (Barton & Capellini, 2011). Such inconsistencies in applicability and explanation raise the question: are we failing to accurately measure and explain brain size and associated traits?

Despite Healy and Rowe’s (2007) warning, studies reporting cross-species correlations between brain size measures and various behavioral and life-history traits continue to accumulate. This is in spite of recent evidence falsifying many of the assumptions listed in §2 (see Montgomery, 2017 for a review). For example, brain size does not scale linearly with body size within (Rubinstein, 1936) or across species (e.g., Fitzpatrick et al., 2012; Montgomery et al., 2013; Montgomery, Capellini, Barton, & Mundy, 2010), brain regions do not scale uniformly with total brain size across species (§4.1; e.g., Barton & Harvey, 2000; Farris & Schulmeister, 2011; Gonzalez-Voyer, Winberg, & Kolm, 2009), brain size does not uniformly scale with neuron number across taxa (§4.1; Herculano-Houzel, Catania, Manger, & Kaas, 2015; Olkowicz et al., 2016), brain size does not necessarily translate into cognitive ability (see §2.3, §3.2, Box 1), and brain size is not consistently related to variables of interest even within species (see §2.2; e.g., there are sex differences with regard to brain size and its relationship with cognition [Kotrschal et al., 2013, 2014]; and fitness and longevity [Logan, Kruuk, Stanley, Thompson, & Clutton-Brock, 2016]). Therefore, a research program that relies on one or more of these assumptions is limited in its ability to make reliable inferences about what brain size measures and what it means when it correlates (or not) with other traits.

### 2.2 Does selection act on brain size?

Attempts to explain variation in brain size often implicitly assume that natural selection acts directly on brain size. In vertebrates this assumption has been given added traction from models exploring how brain development may shape patterns of evolution that place greater emphasis on the conservation of brain architecture. This renders brain size a potent target of selection, in contrast to selective adaptation of particular brain regions (see §4.2). Artificial selection experiments further highlight the capacity for selection to directly act on brain size (e.g., Atchley, 1984; Kotrschal et al., 2013). For example, artificial selection for small and large brain size in guppies (*Poecilia reticulata*) produced a grade-shift in the scaling relationship between brain and body size, resulting in ~15% differences in relative brain size between selection lines (Kotrschal et al., 2013). While the resulting large- and small-brained guppies differed in several traits, including performance in learning tasks (Kotrschal et al., 2013, 2014) and survival (Kotrschal et al., 2015), almost all of these correlations between behavioral performance and brain size were either test context- or sex-dependent (Kotrschal et al., 2013, 2014, 2015; van der Bijl, Thyselius, Kotrschal, & Kolm, 2015).

These various trade-offs and sex-specific effects suggest the selection landscape in natural populations may routinely be more complex than under laboratory conditions. Several recent studies of variation in brain composition among closely related populations or species that are isolated by habitat reveal heritable divergence in particular brain components rather than overall size (Gonda, Herczeg, & Merilä, 2011; Montgomery & Merrill, 2017; Park & Bell, 2010). Indeed, a recent analysis of brain morphology in wild guppies suggests selection may frequently favor changes in the size of specific brain regions, although in this case a role for plasticity has not been ruled out (Kotrschal, Deacon, Magurran, & Kolm, 2017). Focusing solely on overall brain size, as in the artificial selection experiments, might mask the co-occurring changes within the brain that underlie the observed differences in behavior. Accordingly, adaptive responses to ecological change may involve alterations in specific components of neural systems, presumably in response to selection on particular behaviors. This latter distinction is important. It is unlikely that selection ever acts ‘on’ any neuroanatomical trait because what selection ‘sees’ is variation in the phenotypes produced by neural systems (i.e., behavior), and the energetic and physiological costs associated with their production.

Understanding how brain size relates to selection for behavioral complexity or cognition is therefore a two-step process. First, we must understand how behavioral variation emerges from variation in neural systems. Second, we must understand how this variation in neural systems relates to overall brain size. Currently, our ability to take these steps is limited by a paucity of well understood examples of behavioral variation in natural populations. However, existing examples provide some insight into the limitations of total brain size as a unitary trait. Recent studies of the proximate basis of schooling behavior in fish (Greenwood, Wark, Yoshida, & Peichel, 2013; Kowalko et al., 2013), and burrowing (Weber, Peterson, & Hoekstra, 2013) and parental behaviors in *Peromyscus* mice (Bendesky et al., 2017), suggest outwardly unitary ‘behaviors’ may often be composites of genetically discrete behavioral phenotypes whose variation is determined by independent neural mechanisms. For example, in a recent study of parental care in *Peromyscus*, Bendesky and colleagues (2017) used a quantitative genetics approach to map variation in the propensity to care between two species. They revealed a highly modular genetic architecture, with some loci affecting general care behavior and other loci affecting specific traits such as nest building, and a high propensity for sex-specific effects. Variation in specific traits such as nest building in males can be linked with particular brain regions, in this case, the hypothalamus.

The role of *FOXP2*, a transcription factor, in language development and evolution provides a second informative example. *FOXP2* is generally highly conserved across mammals, but has two human-specific amino acid substitutions which were likely fixed by positive selection (Enard et al., 2002). Disruption of this gene in humans severely impacts language acquisition (Lai, Fisher, Hurst, Vargha-Khadem, & Monaco, 2001), suggesting it plays a key role in vocal learning. Insertion of the human version of the protein into the mouse genome affects the development of particular cell types in the basal ganglia without gross effects on brain size or morphology (Enard et al., 2009), yet leads to improved performance on certain learning tasks and may have a broader role in motor learning (Schreiweis et al., 2014).

These examples illustrate how variation in behaviors that are considered by many comparative studies to be correlated with whole brain size may in fact arise from localized changes in brain development that do not affect total size. This may be the kind of incremental variation selection plays with over small evolutionary time scales, and it is reasonable to assume that the accumulation of this kind of change makes a significant contribution to species differences in total brain size. While there is some evidence that genetic pleiotropy (i.e., genetic variation in loci that cause phenotypic variation in multiple traits) can drive shifts in multiple behaviors, in many cases selection may be able to shape specific behavioral traits independently of other behaviors. Global measures of brain size and cognition both suffer from a lack of support for the underlying assumption that the correlated variation in their component parts stems from a shared proximate basis.

### 2.3 Assumptions and limitations about what brains mean for cognition

While ongoing efforts attempt to discover and characterize specific links between particular behavioral and cognitive traits and brain size, this work exists alongside a highly visible thread running through the literature that takes brain size itself as a proxy for ‘intelligence’ (e.g., Jerison, 1969; Table 1). For example, Jerison (1973, 1985) hypothesized that species showing behaviors assumed to require increased neural processing required the evolution of a larger brain relative to their body size to create ‘extra neurons’ for those seemingly complex behaviors.

**Table 1:**
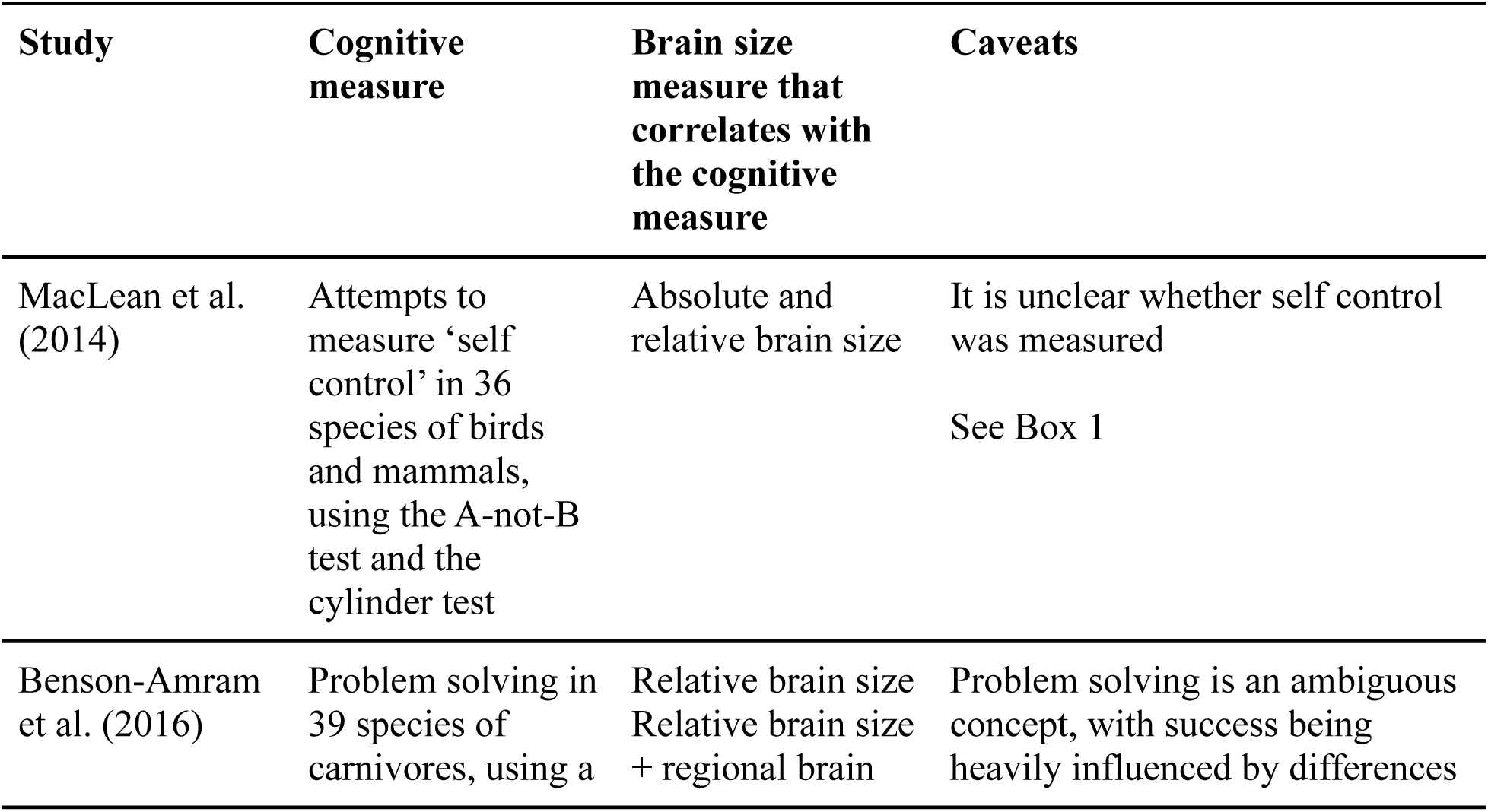
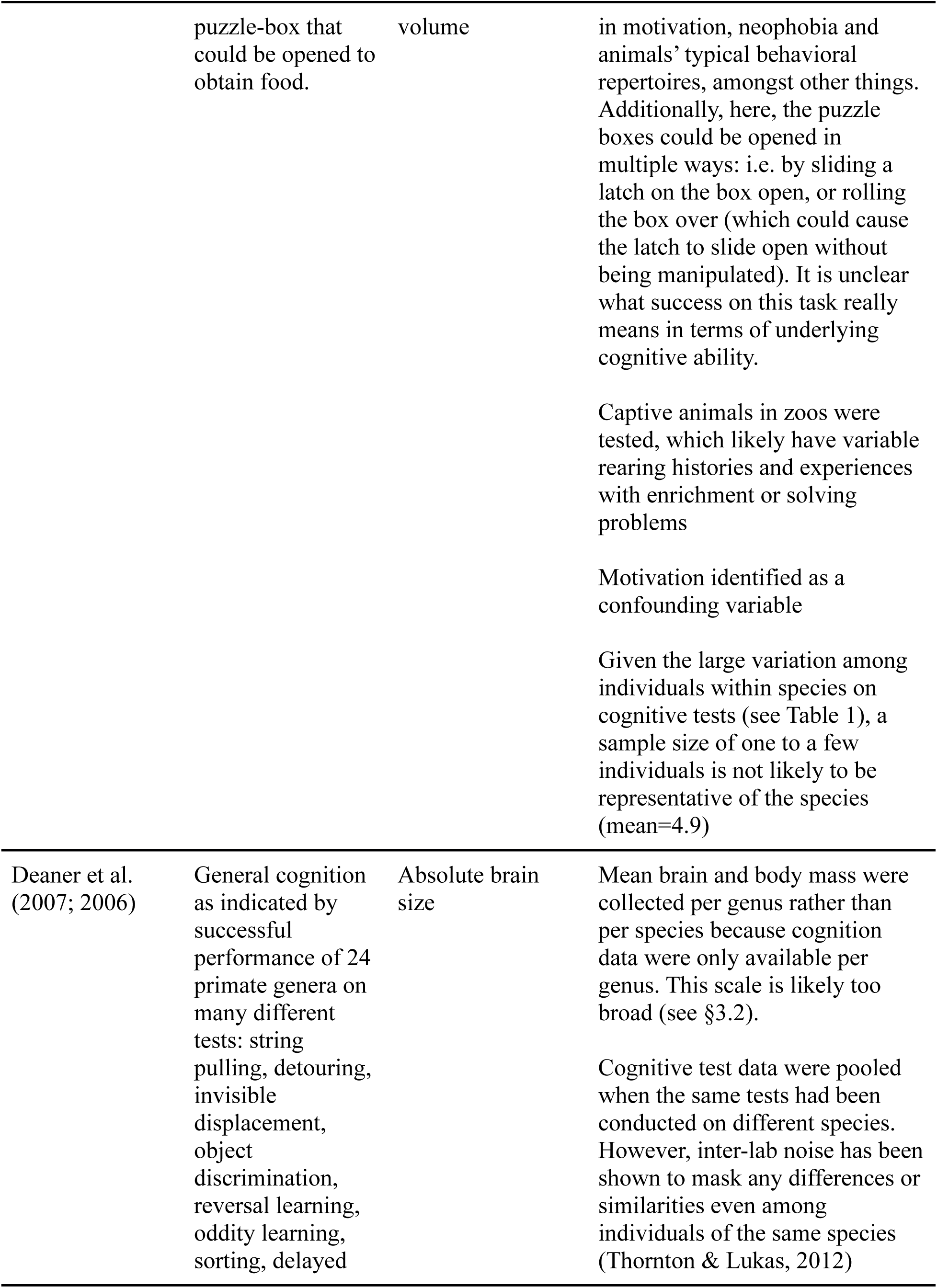
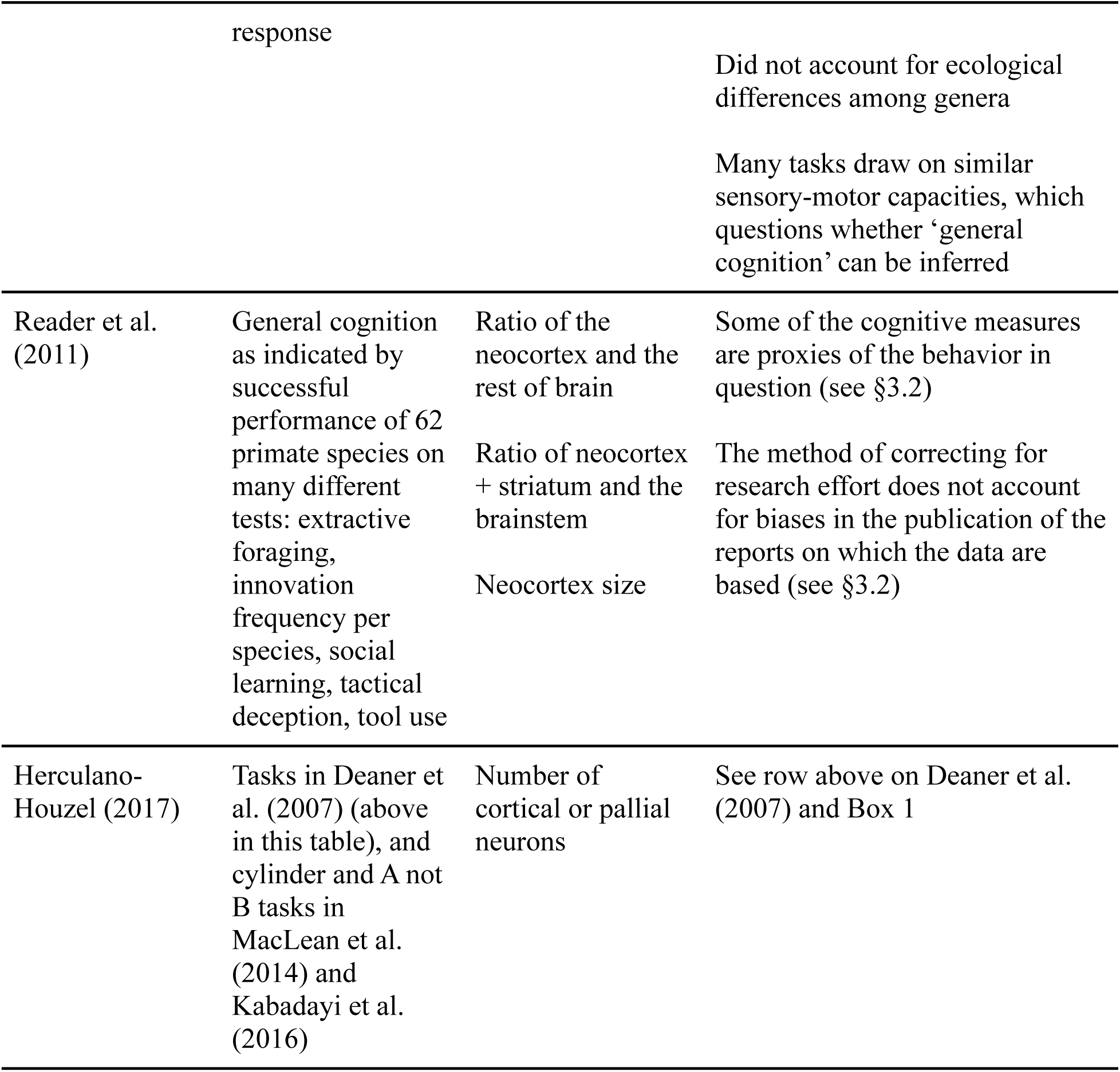
Examples of cross-species comparisons that link cognition and brain size, and a description of the caveats about the ability to draw inferences due to the limitations involved in measuring both traits.

In discussing indicators of ‘cognition’, we first need to know when a behavior is ‘cognitive’ or indicative of ‘complex cognitive abilities’ (sometimes referred to as ‘intelligence’ and often invoking the term ‘behavioral flexibility’ [Mikhalevich, Powell, & Logan, 2017]; Table 2). This is problematic because these terms are not defined well enough to test empirically or even to properly operationalize, and therefore cannot be measured in a systematic way. Appeals to ‘neural processing’ likewise suffer from ill definition and an inability for accurate quantification in most contexts. Researchers studying animal behavior tend to avoid using the term ‘intelligence’ due to its anthropocentric connotations, and instead often adopt Shettleworth’s definition of cognition as “the mechanisms by which animals acquire, process, store, and act on information from the environment. These include perception, learning, memory, and decision-making” (Shettleworth, 2010, p. 4). However, this all-encompassing definition still does not allow us to answer basic questions about the proximate machinery underlying ‘cognitive’ traits: Is a behavior more ‘cognitively complex’ if it engages more neurons, or certain networks of neurons, or neurons only in particular brain regions that are responsible for learning and memory? Or should we think of neural processing in dynamic terms, such as the ‘flexibility’ of neurons to abandon old connections and form new ones as task demands change? Is behavior only considered to rely on complex cognition if it is flexible? There are no clear answers to these questions because data are greatly lacking.

**Table 2:** Examples of experiments attempting to test cognition. We note that there may be additional confounds in such studies that are likely to have affected test performance; however, these cannot be ruled out until explicitly quantified and taken into account in analyses (see also Macphail, 1982).

| Cognitive Tests | Attempting to test | Assumed cognitive measures might be confounded by | Examples of studies that investigate or review these cognitive tests |
| --- | --- | --- | --- |
| String pulling | Insight<br>Learning speed<br>Means-end understanding | Responses to perceptual-motor feedback<br>Motivation to obtain reward<br>Age<br>Attention<br>Rearing effects<br>Dexterity<br>Object permanence<br>Exploration<br>Neophobia/neophilia<br>Side biases<br>Visual acuity<br>Salience of the stimuli | Jacobs & Osvath (2015) |
| Aesop’s Fable tube tests | Physical cognition<br>Causal reasoning | Learning speed<br>Motivation to obtain reward<br>Object biases | Jelbert et al. (2014; 2015), Logan et al. |
|  |  | Responses to perceptual-motor feedback<br>Salience of the stimuli | (2014), Miller<br>et al. (2016) |
| Cylinder<br>and A not B<br>tasks | Self control /<br>inhibition | Neophobia/neophilia<br>Exploration<br>Dexterity<br>Perseveration<br>Salience of the stimuli<br>Visual acuity/tracking<br>(see Box 1) | MacLean et al.<br>(2014), Jelbert<br>et al. (2016) |
| Puzzle-<br>boxes | Innovativeness<br>Problem solving<br>Creativity | Neophobia/neophilia<br>Exploration<br>Dexterity<br>Motivation to obtain reward<br>Behavioral repertoire size<br>Perseveration<br>Operant conditioning (likelihood of<br>repeating actions that led to obtaining<br>reward)<br>Salience of the stimuli<br>Age<br>Dominance rank<br>Sex | Benson-Amram<br>& Holekamp<br>(2012),<br>Boogert et al.<br>(2008),<br>Thornton &<br>Samson (2012) |
| Reversal<br>learning | Learning speed<br>Behavioral<br>flexibility | Neophobia/neophilia<br>Exploration<br>Perseveration<br>Learning speed<br>Motivation to obtain reward<br>Body condition<br>Age<br>Sex<br>Reproductive hormonal state<br>Habituation to captivity<br>Salience of the stimuli | Boogert et al.<br>(2011; 2010),<br>O'Hara et al.<br>(2015) |
| Trap-tube<br>tasks | Tool-use<br>Physical<br>cognition<br>Causal reasoning | Operant conditioning<br>Learning speed<br>Dexterity<br>Neophobia<br>Motivation to obtain the reward<br>Motivation to avoid incorrect responses<br>Inhibitory control<br>Salience of the stimuli | Seed et al.<br>(2006),<br>Mulcahy & Call<br>(2006) |

Indeed, it is nearly impossible to determine which behaviors require increased neural processing when they are observed in isolation from real-time brain activity. Creative studies using imaging technology can now measure behavior and brain activity at the same time, but only in species that can be trained to remain stationary in an fMRI scanner (e.g., dogs: Andics et al., 2016; pigeons: De Groof et al., 2013; see also Mars et al., 2014). However, without *a priori* predictions about which neural measures indicate complex cognition, this will remain a process o f *post hoc* explanations and goal-post moving based on anthropocentric biases about which species should be ‘intelligent’ (see Mikhalevich et al., 2017).

Burgeoning research in artificial intelligence and machine learning suggests the correlation between raw computing power (‘brain size’) and ‘intelligence’ is unlikely to be straightforward. For example, a machine-learning algorithm designed to solve a specific task may indeed get a performance boost from a ‘bigger brain’ (i.e., utilizing more hardware, for example, when playing Go [Silver et al., 2016]). However, algorithmic improvements that create more efficient ways of forming ‘neuronal’ connections based on input data may account for even greater performance or speed improvements given fixed hardware. The effective utilization of hardware resources is itself an active research field within machine learning (e.g., Nair et al., 2015), hinting that a ‘bigger brain’ does not straightforwardly translate into greater speed or better performance.

Theoretical reflection within the field of artificial intelligence has provided alternative definitions of intelligence that highlight the difficulties faced by cognitive ethologists. For example, Legg and Hutter (2007) aim to provide a universal definition that could apply to machine intelligence as well as human and non-human animal intelligence. Informally, their definition suggests: “*Intelligence measures an agent’s ability to achieve goals in a wide range of environments*”. Following Legg and Hutter’s definition (without committing to whether theirs is definitive) clarifies several difficulties with the current approach to evaluating intelligence in non-human animals, and subsequently our ability to relate it to brain size. More specifically:

1. Intelligence is goal-dependent. A behavior, no matter how complex, cannot be counted as intelligent if it does not serve a clear goal. Yet, interpreting goal-orientation in non-humans is inherently difficult, even under strict experimental conditions.
2. Intelligence is environment-dependent. Problematically, behavioral features often associated with complex cognition such as innovation, planning and tool use may have varying degrees of availability or relevance in different environments, which may affect whether they are displayed or not, irrespective of the organism’s ability to display them.
3. Intelligence of an organism is displayed across a range of environments. The few experimental setups usually used to quantify ‘intelligence’ in captive animals may therefore be minimally informative; instead, the ability of an organism to achieve its goals should be evaluated across the range of environments it is likely to encounter within its lifetime.

Regardless of the validity of the definition, these three features – goal orientedness, environment dependency and utility across heterogeneous conditions – highlight the practical limitations of assessing cognition in animals. The focus on utility further illustrates why selection may favor ‘simple’ behavioral solutions to a task, or why the expression of simple behavior does not preclude the ability of an organism to identify and carry out more complex solutions in alternative contexts. If cognition is something akin to problem-solving capacity, then we should develop measures that pay careful attention to the **range** of problems animals face in their **natural environments**, rather than transferring proxies of intelligence in humans that are relevant to the problems humans face in human environments (see Box 2 for an example).

Nevertheless, many comparative studies do find associations between gross measures of brain size and broadly descriptive behaviors. In what follows, we focus on two factors that explain when and why the results of such comparative studies should be treated with caution: biological heterogeneity, and statistical noise and interference.

## 3) Why do these limitations of brain-behavior comparative studies arise? Noise and interference

The lack of consistency in results from comparative studies (see Healy & Rowe, 2007 for an overview) strongly suggests some underlying variability in the relationship between brain size and complex cognition. In attempting to understand the properties of a particular system, it is useful to distinguish between *noise* (exogenous source) and *interference* (endogenous source; Currie & Walsh, n.d. in review) as distinct kinds of confounds in brain-behavior correlations. *Noise* limits our ability to accurately determine and measure the co-evolving brain structures and cognitive abilities (this section), while interference results from the endogenous features of systems interacting with one another (section 4). One major factor shaping limitations of comparative studies of brain size and cognition is the influence of noise and the numerous covariates influencing the reliability and power to detect true associations. Noise results from exogeneous factors that undermine our capacity to extrapolate across data-points. Measurement error is an inevitable source of noise in these studies because behavior is noisy (§3.1), the behaviors observed might not directly reflect single specific cognitive abilities (§3.2), and the feasibility of obtaining brain measures differs across species, thus limiting comparison (§3.3). Reducing noise requires different experimental and analytical approaches.

### 3.1 Measuring behavior is noisy because behavior is noisy

Animal behavior depends on the integration of internal motivational states and external environmental cues. Although many behaviors are largely stereotyped, the kinds of behavioral traits routinely studied in comparative studies of cognition are not. The expression of behaviors that we might interpret as ‘social cognition’ (such as theory of mind) or ‘physical cognition’ (such as tool use) depend on an individual’s internal state and perception of the external environment, factors that are not readily assessed. This introduces a degree of stochasticity in an animal’s behavioral expression and noise in our behavioral measurements.

For example, Japanese macaques (*Macaca fuscata*) provide a famous case of innovation. In this case, two novel behaviors involving washing sweet potatoes before eating them and separating grains from dirt by throwing them in the water, were innovated by a single female (called Imo) and spread through a wild population via social transmission (cf. Allritz, Tennie, & Call, 2013; Kawai, 1965). At a population level, the high rate of social transmission may be impressive, but at an individual level does the innovativeness of Imo suggest some neuroanatomical variation that supports more complex cognition and increased innovation? Imo’s brain may be no more innovative than her peers: she may simply have been in the right place at the right time or more receptive to reward stimuli. Whether inferred behavioral categories such as innovation reflect population-level variation in cognition is therefore unclear. The assumption that innovation and behavioral flexibility reflect similar cognitive processes is extrapolated from anthropocentric concepts and experiences of innovation, but this clearly requires empirical validation.

Developments in artificial intelligence and machine learning, especially in the field of reinforcement learning, suggest one reason to be wary of the possible conclusion that Imo’s innovative ability is due to neuroanatomical variation at the intraspecies level: high levels of task performance can be achieved by systems that combine trial-and-error learning with feedback on their performance in the form of a reward (similar to reward-based associative learning). As reported by Mnih and colleagues (2015), a system trained from raw pixel inputs and reinforced using an environment-provided performance metric (game score) was able to achieve human-level performance in Atari game play by iteratively searching for the patterns that maximize game score. Silver and colleagues (2016) report another achievement of artificial intelligence, namely of human-level performance on the game of Go, with gameplay that has been described as ‘creative’ and ‘innovative’, by the artificial agent first learning to predict expert moves (supervised learning) and then by improving performance through self-play (reinforcement learning). These engineering achievements suggest that the combination of chance, feedback, and repeated iterations (possibly over generations) could yield the same behavioral performance by artificial intelligence, at least in narrow domains, as organisms associated with having ‘complex cognition’.

### 3.2 Measuring cognition through behavior is noisy because we use unvalidated proxies

Cognition is unobservable and must be inferred from behavior (Box 1). Many of the 50+ traits that have been correlated with brain size across species (Healy & Rowe, 2007) are proxy measures of the actual trait of interest (e.g., the number of novel foraging innovations at the species level is a proxy for individual-level behavioral flexibility). This would not be a problem if proxies were validated by directly testing the link between the trait of interest and its correlational or causal relationship with its proxy within a population. However, the proxies used are generally not validated, which contributes noise and uncertainty about what the correlations, or lack thereof, between these trait-proxies and brain size actually mean. An in-depth discussion of innovation illustrates why and how unvalidated proxies are an issue for this field.

The hypothetical link between innovation frequency per species and their relative brain size was originally proposed by Wyles and colleagues (1983). Lefebvre and colleagues (1997) operationalized the term *innovation* to make it measurable and comparable across bird species, defining it as the number of novel food items eaten and the number of novel foraging techniques used per species as anecdotally reported in the literature (see also Overington, Morand-Ferron, Boogert, & Lefebvre, 2009). Innovation is assumed to represent a species’ ability to modify its behavior in response to a change in its environment, and is therefore a trait-proxy for *behavioral flexibility* (e.g., Overington et al., 2009; Reader & Laland, 2002; Sol & Lefebvre, 2000; Sol, Timmermans, & Lefebvre, 2002). Behavioral flexibility is defined here as modifying behavior in response to changes in the environment based on learning from previous experience (Mikhalevich et al., 2017; Swaddle, 2016). Two challenges emerge from this conceptualization: i) how to measure innovation, and ii) how to validate that innovation frequency per species really is an accurate reflection of behavioral flexibility.

It is unclear how to calculate innovation frequency per species, or what its biological significance is to the species in question. For example, Logan (unpublished data) tried to follow standard methods (from Lefebvre et al., 1997; Overington et al., 2009) to quantify the number of innovations in New Caledonian crows (*Corvus moneduloides*). Innovations were extracted from anecdotal reports “if authors used terms like ‘opportunistic’, ‘novel’, ‘first description’, ‘unusual’, ‘not noted before’, and ‘no previous mention in the literature’” (Lefebvre et al., 1997, pp. 550–551). Technical innovations were also extracted from anecdotal reports and defined as falling into one of these categories: “novel technique, novel technique in an anthropogenic context, novel parasitic behaviour, novel commensal behaviour, novel mutualistic behaviour, novel proto-tool behaviour, novel true tool behaviour and novel caching behaviour” (Overington et al., 2009, p. 1002). Logan found two food-type innovations and 10 technical innovations. However, it was unclear how distinct each technical innovation was (e.g., three involve manipulation of *Pandanus* leaves). It was also unclear whether tools that were used in a similar way, but made of different materials should count as separate innovations or the same innovation (e.g., using non-stick materials in a stick-like manner). Finally, and most importantly, it became clear that these innovations were only novel or unusual to the humans who saw crows performing these behaviors; these behaviors are commonly performed by New Caledonian crows across their natural habitat (e.g., Hunt & Gray, 2002) and are certainly not novel to them, suggesting that innovation frequency databases (e.g., Overington et al., 2009) may contain many similar cases of species-typical behaviors that had gone unnoticed to the human observer. Therefore, it is also unclear what innovation frequency per species means to that species, which further confounds the significance of innovation frequency per species.

Evidence has only recently become available to test the hypothesis that innovation frequency per species is a reliable proxy for behavioral flexibility. A small number of comparative studies have tested individuals from different species that vary in brain size and innovation frequency (both are species-level measures) on the same test of behavioral flexibility. Results showed that innovation frequency per species did not correlate with measures of behavioral flexibility in individuals (Auersperg, Bayern, Gajdon, Huber, & Kacelnik, 2011; Bond, Kamil, & Balda, 2007; Ducatez, Clavel, & Lefebvre, 2015; Jelbert et al., 2015; Logan, 2016a, 2016b; Logan, Harvey, Schlinger, & Rensel, 2016; Logan et al., 2014; Manrique, Völter, & Call, 2013; Reader et al., 2011; Tebbich, Sterelny, & Teschke, 2010) or with species level estimates of brain size (Cnotka, Güntürkün, Rehkämper, Gray, & Hunt, 2008; Ducatez et al., 2015; Emery & Clayton, 2004; Isler et al., 2008; Iwaniuk & Nelson, 2003; Pravosudov & de Kort, 2006) in predictable ways. The absence of consistent associations between intraspecific measures of behavioral flexibility and species-level measures of innovation and brain size erodes the logical basis of comparative studies across species. If behavioral flexibility is to be considered a marker of cognitive ability, it should be measured directly in individuals of each species rather than using unsupported species-level proxies such as the reported frequency of innovation. The continued use of innovation frequency is due solely to convenience and data availability. Although the first comparative studies using this metric provided promising glimpses into brain evolution, the time has now come to descend to the within-species level to understand the proximate origins of individual variation in behavioral flexibility. More generally, despite a lack of validation that they accurately reflect the trait of interest, proxies of behavioral traits are pervasive in the comparative brain size literature and introduce unknown amounts of exogenous noise into cross-species analyses. This noise may generate spurious results, masking ‘true’ patterns in the data and impeding their interpretation.

### 3.3 Measuring brain size is noisy because it is more difficult than it seems

If cross-species correlations indicate relationships that are actually present, these associations should persist within species if we assume a direct relationship between brain size and behavior in a given task. However, these associations often do not persist at the intraspecies level, which may be due to extensive measurement errors in quantifying brain size within species, or to the confounding effects of variation in a brain trait being attributable to multiple functions. Most work on brain evolution has focused on overall brain size or changes in large regions of the brain, such as the forebrain and the cerebellum (see review in Healy & Rowe, 2007; see also Herculano-Houzel, 2012; Reader et al., 2011). However, volumetric measurements are particularly noisy. We use primate brain data to illustrate the difficulties involved in obtaining, preserving, and measuring brain volumes.

More is known about brain anatomy in primates than in other orders, yet volumetric measurements of specific brain regions in this group are only available for a few species, and some of these measurements are pooled from only a few individuals per species (Reader & Laland, 2002). This introduces a large amount of noise because a species’ average brain, or brain region, volume might be biased due to sexual dimorphism or other variables that differ across individuals (Montgomery & Mundy, 2013). Information on primate brain size is scarce and primarily comes from captive individuals. Further, access to primate brains is limited to only a few brain collections (Zilles, Amunts, & Smaers, 2011).

Additional complications arise in determining whether it is appropriate to correlate behavioral data from wild individuals with morphological data (e.g., brain size) obtained from captive individuals. Studies comparing the morphology of wild and captive animals have shown that rearing conditions may influence body composition (e.g., skull shape, brain size, digestive tract) after only a few generations (O’regan & Kitchener, 2005). In primates, brain mass is not generally affected by captivity (Isler et al., 2008), but body mass is: some species become heavier, while others become lighter due to inadequate diets (O’regan & Kitchener, 2005).

Furthermore, although brain size might not be affected by captivity, primate populations of the same species that were reared under different captive conditions differ in cortical organization (Bogart, Bennett, Schapiro, Reamer, & Hopkins, 2014). In macaques and humans, there is evidence that individual differences in social network size correlate with amygdala volume and areas related to this structure (Bickart, Wright, Dautoff, Dickerson, & Barrett, 2011; Kanai, Bahrami, Roylance, & Rees, 2011; Sallet et al., 2011). Among individuals of the same species, brain anatomy changes significantly with age (Hopkins, Cantalupo, & Taglialatela, 2007). Choosing individuals with closely matched histories can reduce noise in brain measures that are introduced by individual differences in previous experience, but the noise involved in brain volume measurements is most effectively controlled and minimized by obtaining large sample sizes per species to acquire more reliable species averages. This problem is particularly vexing when combining behavioral data sets from observations in the wild with neuroanatomical data from captive populations.

Data collection methods can also compromise the quality of the data. Many reported brain weights and brain volumes are actually proxies of these measures, obtained instead by calculating endocranial volume from skulls, which are much easier data to collect (e.g., Isler et al., 2008; Iwaniuk & Nelson, 2002). While endocranial volume has been shown to reliably approximate brain volume across species of primates (Isler et al., 2008) and birds (Iwaniuk & Nelson, 2002) and within species of birds (Iwaniuk & Nelson, 2002), this might not always be the case. For example, Ridgway and colleagues (2016) suggest endocranial vascular networks and other peripheral appendages can account for 8–65% of endocranial volume in cetaceans, leading to a consistent overestimation of brain size that is more severe in some species than others.

Because brains are valuable tissues, non-invasive methods such as magnetic resonance imaging (MRI) are preferred for obtaining data on brain anatomy and function. Yet high-resolution, high-quality MRIs from primate brains are difficult to obtain from live individuals. Images obtained using in vivo techniques, where the animal is sedated for a short period of time while scanning their brains, might be more accessible, but image quality and resolution is poorer than in images obtained post-mortem (K. L. Miller et al., 2011). Post-mortem MRIs can have a higher resolution and are therefore more suited to calculating volumes. However, even MRIs are problematic because of other sources of noise that arise from brain extraction methods, including the post-mortem delay between death and extraction and preservation, and the ‘age’ of the preserved brain (i.e., how long a brain has been stored for; Grinberg et al., 2008; K. L. Miller et al., 2011). While post-mortem MRI is the best method available for calculating brain volumes, brain volume in itself is a noisy measure because of its unclear, and usually untested, relationship with other variables of interest (see §3.2).

Given that volumetric brain measurements suffer from many additional sources of noise, it may be more productive to focus instead on non-volumetric features of brain composition (e.g., neuron density, grey matter as a measure of local connectivity, white matter as a measure of long-distance connectivity). For example, transcranial magnetic stimulation is increasingly used in humans to temporarily ‘knock out’ particular brain areas to understand their functionality and relationship with behavior and cognition (e.g., Zatorre, Chen, & Penhune, 2007). This type of (non-volumetric) technique allows one to elucidate causal relationships, which generates data of a much higher quality because it is validated (i.e., not a proxy) and directly connected with the behavior under study, which greatly reduces measurement noise. Ideally, multiple methods should be applied to the same system to determine whether different types of evidence arrive at the same conclusions regarding brain-behavior relationships.

## 4) Why do limitations in brain-behavior comparative studies arise? Evidence of interference

Interference occurs when systems consist of multiple interacting parts whose interactions tend to be complex. A potentially useful way of understanding some critiques of brain size-cognition comparative studies is to consider the ramifications of heterogeneity within and across species in terms of their brain architectures and associated traits (e.g., behavior, cognition, life history; Figure 1). If parts of the brain evolve in concert due to developmental coupling, for instance, then interference from those components makes it difficult to isolate the evolutionary causes of changes in brain size, or any of its components, over time. Similarly, if many ecological and life history traits covary, identifying which factors drive changes in brain size is complicated by autocorrelation between independent variables. Philosophers distinguish *heterogeneity* within and between systems as a useful concept for framing the validity of comparisons (Elliott-Graves, 2016; Matthewson, 2011). Heterogeneity arises as a confounding factor in comparisons among individuals and/or species when the components of a system (e.g., brain structures) differ (§4.1), or when similar components exist but differ in scaling relationships or patterns of connectivity (e.g., neuron density, neural network; §4.2). Treating brain size as a unitary trait assumes either that the brain is a unitary trait or that any signal from a brain-behavior association is sufficient to overpower the influence of heterogeneity on either trait. Comparisons of taxonomically diverse neural systems can identify where similar brain architectures exist and where heterogeneity in brain composition is masked by comparisons of brain size (§4.3). Interference in the form of heterogeneity between systems occurs because of the complex interactions among life-history and ecological factors that shape the co-evolution of cognitive abilities and particular brain measures (§4.4).

**Figure 1:**
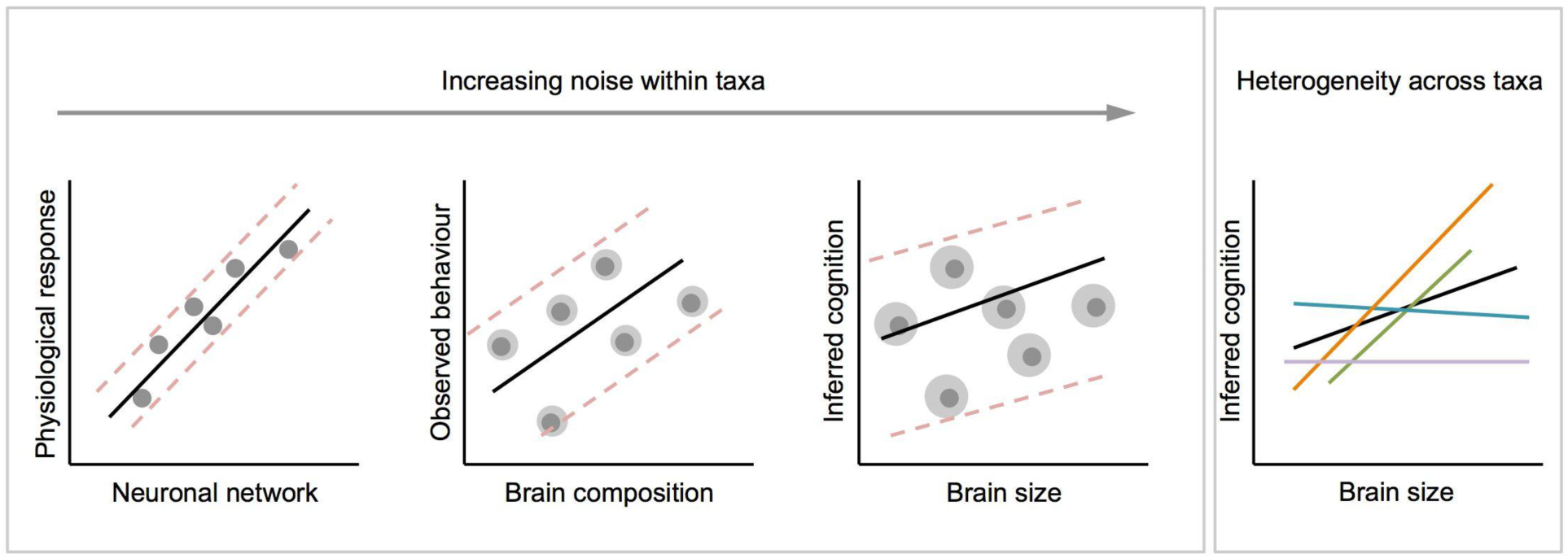
Effects of noise and heterogeneity on brain-behavior correlations as measures of a biological trait (on both axes) become increasingly crude. As measurements move away from direct, quantitative data of primary biological processes both axes become increasingly noisy (as indicated by the grey halos around each data point). The interaction between signal, noise, and heterogeneity may result in contrasting correlations between taxonomic groups (indicated by differently colored lines). When correlations are averaged across these groups the resulting associations may retain little information.

### 4.1 Heterogeneity in brain composition within taxonomic groups: brains that appear similar according to certain measures may actually be different

Broad comparisons across phylogenetically disparate and ancient groups demonstrate how our understanding of the presumed cognitive benefits of large brains is, at best, simplistic. The brain architecture underlying ecologically relevant neural computation will depend on the behavioral requirements of a task, the evolutionary history of the machinery that selection is building on, and the strength of potentially opposing selective forces such as energetic, volumetric, and functional trade-offs and constraints. Even across more closely related species, for example among mammals, heterogeneity between brain structures introduces noise and variation that can complicate brain-behavior relationships.

While some authors argue that the major axis of variation in mammalian brains is overall size (e.g., Clancy, Darlington, & Finlay, 2001; Finlay, Darlington, & Nicastro, 2001), there is ample evidence for variation in brain structure across species caused by brain region-specific selection pressures, so called ‘mosaic brain evolution’ (Barton & Harvey, 2000). Accumulating evidence among major taxonomic groups shows differences in brain composition (e.g., Kaas & Collins, 2001; Workman, Charvet, Clancy, Darlington, & Finlay, 2013). When a behavior generated by a specific brain structure is targeted by selection, the effect on total brain size will depend on the scaling relationship between that brain structure and total brain size. For example, one general trend across mammalian brain evolution is a correlated expansion of the neocortex and cerebellum, which occurs independently of total brain size (Barton, 2012; Whiting & Barton, 2003). These structures share extensive physical connections and are functionally interdependent (Ramnani, 2006), but, while they tend to co-evolve, both have evolved independently in some evolutionary lineages (Barton & Venditti, 2014; Maseko, Spocter, Haagensen, & Manger, 2012). Independent selection pressure on individual brain components such as the neocortex and cerebellum do not have equal effects on overall brain size or measures of encephalization (Figure 2). Neocortex volume scales hyper-allometrically with brain volume (i.e., as brain size increases, the proportion of neocortex tissue increases), while cerebellum volume, and several other major brain components, scale hypo-allometrically with brain volume (Barton, 2012). As a result, increases in neocortex volume have a disproportionate effect on brain volume compared with similar proportionate increases in cerebellum size, largely due to differences in the scaling of neuron density and white matter in the two structures (Barton & Harvey, 2000; Herculano-Houzel, Collins, Wong, & Kaas, 2007). Variations in whole brain size, or measures of brain size relative to body size, such as the encephalization quotient (Jerison, 1973), therefore essentially correspond to variation in neocortex size and mask variation in other brain components, even though the latter may be of great functional significance. The power of the comparative analysis of brain-behavior associations is therefore limited when selection acts on non-cortical structures. Even in vertebrates, mounting evidence suggests this will often be the case. For example, the frequency of tool use in primates (Barton, 2012) and the complexity of nest structure in birds (Z. J. Hall, Street, & Healy, 2013) have been linked with variation in relative cerebellum volume, and spatial memory in birds has been linked with hippocampal volume (e.g., Krebs et al., 1996).

**Figure 2:**
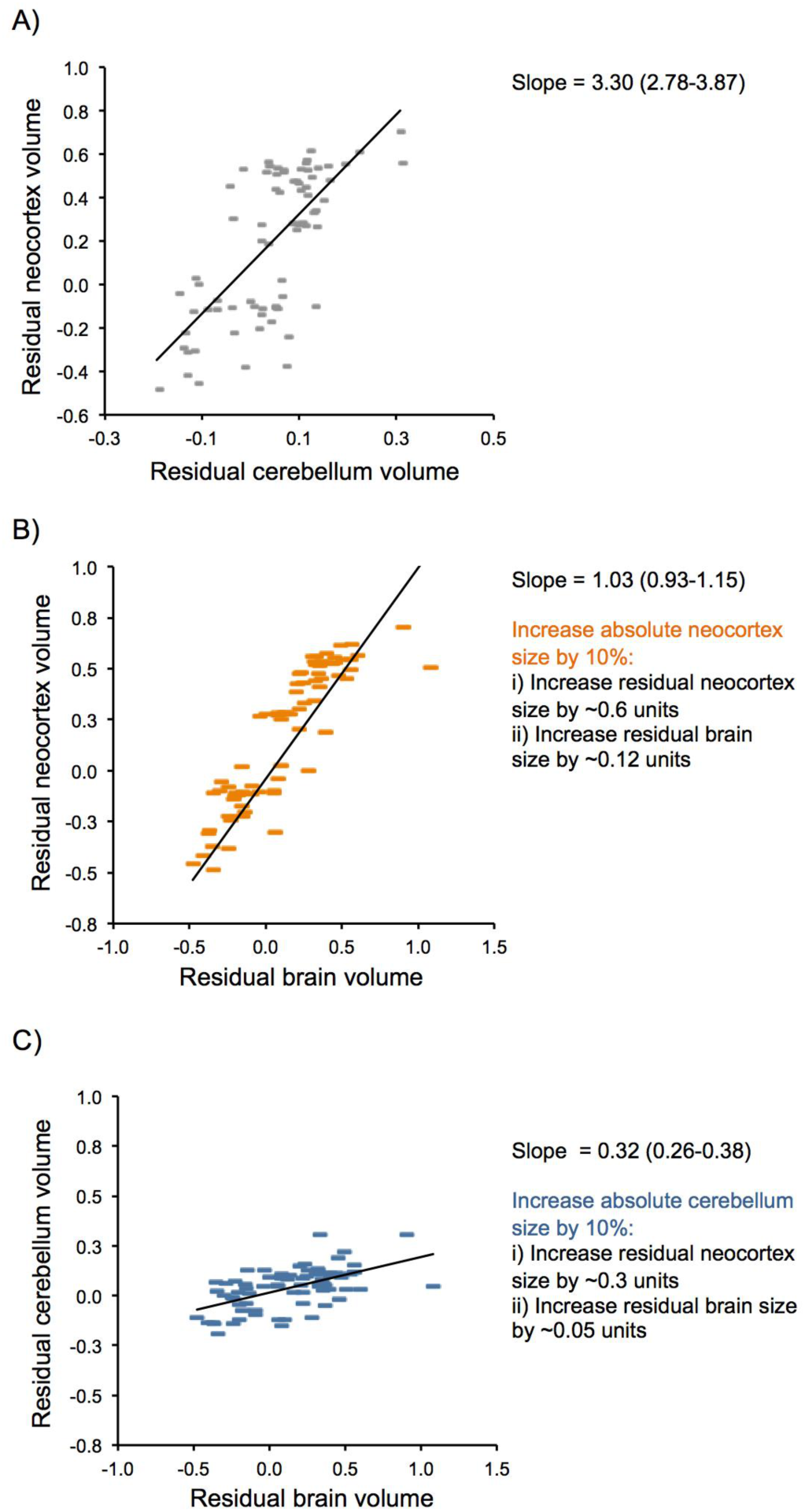
Effects of brain component scaling on the contributions brain regions make to brain expansion. A) the size of the neocortex and cerebellum, once corrected for the size of the rest of the brain, co-evolve with a positive scaling relationship. Both residual size of the neocortex (B) and cerebellum (C), after correcting for the size of the rest of the brain, correlate with the total brain size corrected for body size indicating both components contribute to encephalization. However, the scaling relationships differ, such that any increase in absolute neocortex volume has a greater influence on residual brain size compared to a similar increase in absolute cerebellum volume (see also Barton, 2012).

When comparing brain size across species, further heterogeneity is apparent at the level of the cellular composition of brain structures. Recent data on neuron number in brain regions of birds and mammals have revealed extensive variation across taxonomic groups (Herculano-Houzel et al., 2015). For example, primates have significantly higher neuron densities in the neocortex and cerebellum than other closely related terrestrial mammals, while elephants have substantially higher neuron densities in the cerebellum than other Afrotheria (e.g., golden moles, elephants, and sea cows; Herculano-Houzel et al., 2015), and birds pack similar numbers of neurons as found in primates into volumetrically more restricted brains (Olkowicz et al., 2016). Because neurons and their synaptic connections are the basic computational units of any neural system, if variation in brain, or brain region, volume does not consistently reflect variation in neuron number, then any inference made about the cognitive significance of brain size is largely invalid. To illustrate this effect, averaging across brain regions, a 1 gram brain that follows primate neuron number-brain size scaling rules will contain ~26% more neurons than a brain that follows the glire scaling rules (the clade including rodents; Herculano-Houzel et al., 2015). A 1 gram brain that follows psittacine (parrots) scaling rules will contain ~100% more neurons than a brain that follows the glire scaling rules, and ~58% more than one that follows the primate scaling rules (Olkowicz et al., 2016). Comparing brain size across these taxa thus erroneously assumes that the computational output (based on neuron number) of these hypothetical brains would be equal.

The assumption that brain volume is comparable and meaningful across species is explicitly made in broad phylogenetic studies of cognitive ability (e.g., MacLean et al., 2014; Box 1). Variation in brain structure and cellular composition strongly questions this assumption. The effect of incorporating more fine-grained data, even if they are relatively crude, is apparent in existing studies. For example, in Benson-Amram and colleagues’ (2016) analysis of how performance on a puzzle-box test is associated with brain size across 39 species of mammalian carnivore, the addition of data on volumetric variation in brain structure significantly improved their predictive model compared to one containing only brain volume. In a recent opinion piece, Herculano-Houzel (2017) also argued that (cortical) neuron number outperforms total brain size as a predictor of behavioral performance in self-control tests reported by MacLean and colleagues (2014). The power of brain size as a causative predictor of cognitive performance is therefore apparently vulnerable to the addition of only narrowly more fine-grained data. The sole reasons for the continued focus on brain size as a unitary trait are convenience and data availability. If we are to progress beyond a superficial understanding of brain-behavior correlations, this justification must also be set aside now that new forms of more detailed data are becoming increasingly available on how neuron number and connectivity vary across brain regions and species.

### 4.2 Deep convergence in brain architecture: brains that appear different according to certain measures may actually be homologous

At the broadest taxonomic scale, brain composition is remarkably diverse. For example, comparative studies have traditionally focused on linking learning and memory with arachnid protocerebrums (e.g., Meyer & Idel, 1977; Punzo & Ludwig, 2002), insect mushroom bodies (e.g., Snell-Rood, Papaj, & Gronenberg, 2009), cephalopod vertical lobes (e.g., Grasso & Basil, 2009), the vertebrate pallium (e.g., Jarvis et al., 2005), and mammalian neocortices (e.g., Pawłowskil, Lowen, & Dunbar, 1998). Despite their independent evolution, some research points toward commonalities in the molecular and neural systems that function in heterogeneous brain organizations across animal phyla. A combinatory expression pattern of developmental control genes suggests the deep origin of key learning and memory centers, including in the complex sensory centers and cell types of the mushroom bodies of annelids and arthropods, and the pallium of vertebrates (Tomer, Denes, Tessmar-Raible, & Arendt, 2010). Similarly, Pfenning and colleagues (2014) proposed that vocal learning in birds and humans has evolved via convergent modification of brain pathways and molecular mechanisms. Roth (2013, p. 292) proposed that the centers for learning and memory in insect, octopus, avian, and mammalian brains share a comparable associative network that “bring[s] the most diverse kinds of input into the same data format and [integrates] the respective kinds of information.” These broad comparisons suggest that such brain structures in taxonomically and anatomically diverse animals may share a number of features, including high neuron density, and similar organizations with hierarchical connectivity (G. Roth, 2013). Similarly, the vertebrate basal ganglia and insect central complex have been shown to exhibit a deep homology, sharing similar network organizations, neuromodulators, and developmental expression machineries (Strausfeld & Hirth, 2013). Accordingly, divergent structures may have converged on similar architectures and computational solutions to analogous behavioral challenges (Shigeno, 2017). By simplifying brain measures by focusing only on size, we may miss out on opportunities to study how convergences in behavior and complex neural systems can inform how cognition evolves.

Nevertheless, the heterogeneity identified by these studies may also provide useful variation that can contribute to our understanding of brain and cognitive evolution. For example, if neuronal density can vary independently of volume, why? And how does this impact the functional properties of the pathways that produce complex behaviors associated with cognitive prowess?

#### 4.3 Effects of size-efficient selection

Compared with vertebrates, arthropods have tiny brains and vastly fewer neurons in their nervous systems (Eberhard & Wcislo, 2011), yet many insects and spiders display highly sophisticated motor behaviors, social organizations, and cognitive abilities (Chittka & Niven, 2009; Box 2). For example, insects and spiders exhibit numerical cognition (Cross & Jackson, 2017; Dacke & Srinivasan, 2008; Rodríguez, Briceño, Briceño-Aguilar, & Höbel, 2015), planning (Cross & Jackson, 2016; Tarsitano & Jackson, 1997), selective attention (Jackson & Li, 2004) and working memory (Brown & Sayde, 2013; Cross & Jackson, 2014; Zhang, Bock, Si, Tautz, & Srinivasan, 2005)—all typically studied in vertebrates and considered cognitively demanding (Chittka & Niven, 2009), illustrating that selection has favored highly efficient neuronal systems in these taxa.

Although heterogeneity in brain systems limits the scope of comparative studies of brain size, it also provides an opportunity to understand how selection acts on neural systems, and why selection favors particular solutions over others. One key factor may be the role of size-efficient selection and redundancy in nervous systems. Neurons are energetically expensive cells, and their total cost scales predictably with the size of neural systems (Laughlin, de Ruyter van Steveninck, & Anderson, 1998). Selection must therefore constantly trade-off behavioral performance with energetic and computational efficiency. Exploring how these trade-offs are resolved in real and artificial systems has the capacity to greatly inform why some animals invest in larger brains, while others do not (Burns, Foucaud, & Mery, 2010; Chittka & Niven, 2009; Chittka, Rossiter, Skorupski, & Fernando, 2012; Menzel & Giurfa, 2001).

While an imperfect analogy, researchers’ experience with training artificial neural networks provides an insight into how efficient neural networks can be constructed. Indeed, researchers who aim to create an artificial network that serves as a pattern-learning machine have been largely inspired by the organization of the cerebral cortex in mammals (Mnih et al., 2015). This comparison between artificial networks and cerebral cortex organization was made more notable with recent advances in deep convolutional neural networks (an artificial neural network with a large number of intermediary layers, specialized in identifying patterns in perceptual inputs) such as the deep-Q network (DQN). Beyond mammals, this layer-like organization can also be identified in the brains of for example the common octopus and *Drosophila*, suggesting that a common functionality of information processing patterns may be represented both in artificial and biological neural networks (Shigeno, 2017).

One of the key messages from such research is that training large neural networks is still difficult (Bengio, Simard, & Frasconi, 1994; Glorot & Bengio, 2010; Pascanu, Mikolov, & Bengio, 2013). Even when training is successful, it requires a great deal of time and input data, but more importantly, training too-large a network without the right algorithm often simply fails. In artificial systems, this happens when feedback from the environment is used by the neural network to determine certain flexible values of the computational architecture (e.g., connections between artificial neurons). This problem scales up: greater numbers of flexible values (i.e., network parameters, which grow in tandem with ‘brain size’), require greater amounts of input data and increasingly complex algorithms. Such trade-offs are likely also faced by biological organisms. Thus, in addition to the energetic costs of larger brains, there are also informational costs (i.e., a need for more, better, and/or faster inputs) and computational costs (i.e., efficient ways to use inputs, which may be architecturally difficult for natural selection to find) that limit brain size and may channel the response to selection away from simple increases in the total size of the system or brain.

The hand of size-efficient selection can also be seen in the network architecture of large brains that display a ‘small-world’ topology (Ahn, Jeong, & Kim, 2006; Chen, Hall, & Chklovskii, 2006), which minimizes energetically costly long-range connections in favor of proportionally high local connectivity (Bullmore & Sporns, 2012; Buzsáki, Geisler, Henze, & Wang, 2004; Watts & Strogatz, 1998). Yet, if network architecture is constrained by energetic costs, then what does the evidence of variation in cellular scaling between brain components and across species tell us about how brains evolve?

Variation in the scaling of neuron number with volume likely reflects differences in cell size and patterns of connectivity between neurons. The low neuron density in the neocortex in mammals, compared to that of the cerebellum, reflects the high proportion of the neocortex given over to white matter that consists of mid- to long-range fibres connecting neurons (Ringo, 1991). Variation in the pattern of neuronal connections, and integration between brain regions may help explain variation in cellular scaling. Similar explanations may also apply to scaling differences across taxa, with the high neuronal density of primates being associated with relatively smaller volumes of white matter and connectivity (Ventura-Antunes, Mota, & Herculano-Houzel, 2013). However, these scaling differences could also be driven in part by external influences related to ecology, body size, and morphology. Body size affects many aspects of an animal’s ecology, diet and energy consumption, and physiology (LaBarbera, 1986). It should be no surprise that this may extend to brain composition. For example, the ancestor of extant primates, and most of its descendants, occupied arboreal niches (Cartmill, 1972) and had arboreal locomotor strategies that constrain body size and favor a low center of mass; a strategy that is likely inconsistent with volumetrically expensive modes of brain expansion. Selection pressures that favored the evolution of increased neuron number may therefore have been constrained by the physical demands of occupying an arboreal niche, resulting in changes in neural development that were associated with increased neuron density. Similar, but stronger, selection regimes may also explain the extremely high neuron densities in bird brains (Olkowicz et al., 2016). Conversely, the much lower neuron densities of cetaceans (Eriksen & Pakkenberg, 2007) would be consistent with brain evolution along a trajectory relatively free of size or locomotor constraints.

The expectation that brain size should be a simple predictor of cognitive performance ignores the effect of size-related selection pressures (Chittka & Niven, 2009; Chittka et al., 2012). Size-efficiency is most obvious when considering brain function in small invertebrates, but mounting evidence suggests that the same principles may apply even among vertebrates occupying distinct ecological niches that define the range of permissible body sizes and architectures (Olkowicz et al., 2016). Body size is regularly used as a ‘size-correction factor’ on the assumption that residual brain size is more cognitively relevant, but variation in body size itself reflects the presence of wider ecological and physical selection pressures that may render brain composition and function more divergent than size alone (Fitzpatrick et al., 2012; Montgomery et al., 2013, 2010).

### 4.4 Correlations suffer from interference

Problems of noise are compounded by interference from the complex relationships between many behavioral and anatomical traits. This ‘interference’ not only influences our ability to determine whether a mechanistic link exists between specific brain measures and a certain behavior or cognitive ability, but also in determining their functional link and their adaptive evolutionary history. The comparative study of different species can provide insights into how differences in behavior link with differences in brains (Harvey & Pagel, 1991), and phylogenetic comparisons have been the most widely used approach to test hypotheses about adaptation (see §2.2). However, in addition to relying on unvalidated proxies, adaptive stories are frequently based on correlations. It is therefore necessary to identify potential interference from unmeasured variables to gather evidence for causation before we can accept such adaptive accounts as accurate.

There are four main ways in which interference limits the potential to interpret whether correlations represent adaptations. First, any association between differences in brain measures and behavior might not be direct, but caused by interfering factors. For example, increases in brain size and group size both appear to occur in species that eat foods with high nutritional value, therefore the correlation between brain size and group size might be the result of noise from dietary changes (Clutton-Brock & Harvey, 1980; DeCasien, Williams, & Higham, 2017). Second, even if population studies indicate that a measure of brain size and a behavior are directly linked, comparisons across species cannot immediately reveal the causal direction of the association. For example, an association between increased brain size and decreased risk of predation might result from large-brained species being better able to avoid predation (Kotrschal et al., 2015), or from species with low predation pressure having the opportunity to invest additional resources into brain growth (Walsh, Broyles, Beston, & Munch, 2016). Third, external factors frequently mediate the expression of any link across taxonomic groups. For example, switching to a frugivorous diet might lead to selection on olfactory ability in nocturnal species and visual abilities in diurnal species, resulting in independent episodes of brain expansion driven by selection on distinct sensory modalities and brain components (Barton, Purvis & Harvey, 1995). Fourth, any current link between brain size and behavior might be the product of co-option, after the initial evolution of that brain aspect, rather than the driving selection pressure itself. For example, abilities such as object permanence (i.e., the ability to recall the presence of an out-of-sight object) might have been selected because individuals need to remember the spatial position and temporal availability of food sources in their home range, but it could subsequently be used to distinguish social neighbors from strangers (Barton, 1998). Similarly, selection for improved visual acuity in foraging primates may have later been co-opted to serve in individual recognition and social cognition (Barton, 1998). Although some attempts have been made to tease apart these relationships using path analysis (Dunbar & Shultz, 2007b), this approach still suffers from the effects of co-linearity among variables and does not provide a mechanistic understanding of causative relationships (Petraitis, Dunham, & Niewiarowski, 1996). Recent advances provide some ways to overcome these limitations in the comparative approach (see §5.4), but, as previous authors have pointed out (Garland, Bennett, & Rezende, 2005; Gonzalez-Voyer & Hardenberg, 2014; Harvey & Pagel, 1991), interference fundamentally limits our ability to determine past evolutionary processes based on simple observations of species alive today.

These effects are likely to be particularly influential in the small data sets that characterize many comparative analyses of cognition and brain measures, due to the difficulty in obtaining data. With small data sets, correlations are unlikely to be stable, unless the effect size is large, or noise and interference are low (Schönbrodt & Perugini, 2013). In the vast majority of studies, accuracy and sample size are directly traded-off against one another due to logistical and cost constraints. While this is inevitable, studies aiming for broad phylogenetic comparisons by relying on crude proxies of cognition supposedly measurable across very divergent taxonomic groups may be futile. Any trade-off that reduces accuracy to increase taxonomic breadth risks relying on invalid measures, resulting in unstable and potentially meaningless correlations. Comparisons across large, diverse taxonomic groups can be helpful to identify and describe patterns of variation; however, key insights into the evolutionary history of traits and their associations will be gained by incorporating detailed population studies (see §5.2). As neuronanatomical, behavioral, and statistical tools become increasingly comprehensive and sophisticated, the solutions to these issues will be reachable in the near future.

## 5) Beyond brain size

### 5.1 Matching the right tool with the right question

In the last two sections we emphasized how heterogeneity in brain composition and behavior/cognition, and the subsequent noise this generates can influence our attempts to measure the relationships between these variables. We think these issues motivate turning from coarse-grained, ‘taxon-neutral’ (or hominid-inspired) measures to more local, taxon-specific studies. This is not to say that heterogeneity on its own undermines existing ‘monolithic’ narratives of brain size and behavioral complexity. Rather, these narratives ignore the complexity of links between brain morphology, body morphology, and behavior, and often abstract away from the important ecological and evolutionary drivers of complex behavior that we are trying to understand. We therefore argue against privileging anthropocentric measures or criteria. Instead, we urge a recognition of the multi-dimensional and multi-leveled structure of brains, as well as the disparate and varied ways that brains evolve—in conjunction with bodies, and in response to specific environments—to produce complex behavior. Understanding how brains evolve in response to selection on behavioral complexity or cognition is therefore a two-step process. First, we must understand how behavioral variation emerges from variation in neural systems. Second, we must understand how brains change across species and how this might relate to differences in adaptive regimes.

Discovering and probing correlations between properties of brains and behavioral features can be part of a powerful comparative approach, but we should be wary of reification: mistaking an operationalized target of measurement with a ‘real’ object (Whitehead, 1925). There is a difference between something being measurable and it being causally meaningful. We, and others (e.g., Chittka et al., 2012; Healy & Rowe, 2007), have questioned whether coarsegrained, cross-taxa measurements, such as the encephalization quotient, pick out relations that are in fact explanatory of the evolutionary and developmental relationships between brain, cognition, and behavior across lineages. In fact, similar arguments have been made since scientists first started comparing brain measures across species (Snell, 1892). Instead, we argue for an increased focus on a ‘bottom-up’ approach that begins with i) measurements of features that can be validated within particular taxa in *ecologically relevant* experimental contexts, before ii) testing the evolutionary variability in the relationships between brains and behavior across related species. This will help avoid reification by starting with intraspecific, experimentally verifiable causal connections. The first task involves probing how various taxa respond behaviorally to their environments and other stimuli and determining whether those properties correlate with brain measures in revealing ways. These brain measures will frequently be more fine-grained than brain size, concerning particular neuroanatomical and/or neurophysiological features. The second task involves the construction and testing of hypotheses about the ancestral and evolutionary relationships between those taxa, enabling us to expand to broader categories and correlations in a careful, piecemeal fashion. We expect the results of these two tasks to relate in dynamic ways: considerations of evolutionary scenarios are likely to highlight new kinds of experimental tests and hypotheses in local contexts; and these scenarios will depend crucially on information about local taxa.

### 5.2 Bottom-up vs. top-down

The top-down approach uses cross-species correlations between brain measures and a trait of interest and can be useful for generating hypotheses. However, while these are important for motivating research into the links between brains and behavior, we argue that specific hypotheses should then be tested at the within-species level: from the bottom up. The bottom-up approach involves directly testing behavior and cognition in individuals to determine how they relate to brain measures in these particular individuals of a particular species (ideally measured at the same time as behavior/cognition) to build validated, causative correlations (Chittka et al., 2012). When sufficient data on individuals from a variety of species have accumulated, phylogenetic meta-analyses can be conducted to test whether consistent patterns emerge and hold within and across species (see §5.4; Table 3). Correlations within contemporary populations can tell us whether processes are homologous or analogous across species and show the limits of which processes are likely to occur.

The contrast between top-down and bottom-up approaches is often presented as a difference in terms of investigating the ultimate (top-down looking at adaptations and fitness) versus proximate (bottom-up looking at mechanisms and development) reasons for the evolution of a trait (Laland, Sterelny, Odling-Smee, Hoppitt, & Uller, 2011; Scott-Phillips, Dickins, & West, 2011). However, the approach we suggest does not necessarily make this potentially problematic distinction (Beatty, 1994; Calcott, 2013; Cauchoix & Chaine, 2016; Laland, Odling-Smee, Hoppitt, & Uller, 2013). Our main argument for a bottom-up approach is to encourage researchers to have a clear understanding of what they are investigating rather than to rely on proxies. Detailed individual-based studies can reveal not only which brain measures are involved in a particular cognitive ability or behavior, but also provide important insights into the ecological correlates and fitness consequences of variation in particular brain measures (Table 3). Further, starting from behaviors in particular species makes ensuring ecological, developmental, and evolutionary relevance significantly more straightforward: it is a strategy for both avoiding reification, and for being sensitive to the heterogeneity of both brains and behavior.

**Table 3:**
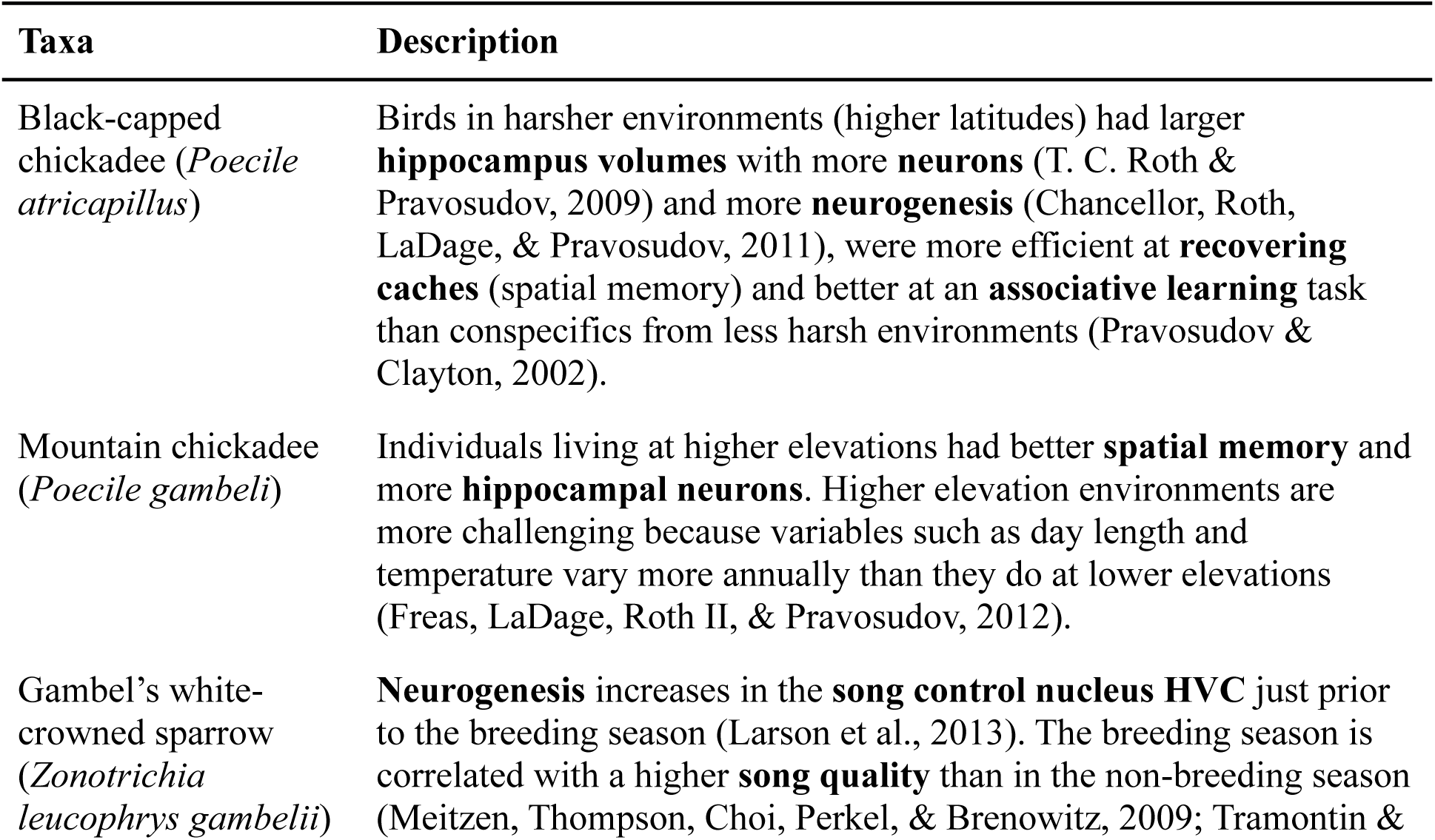
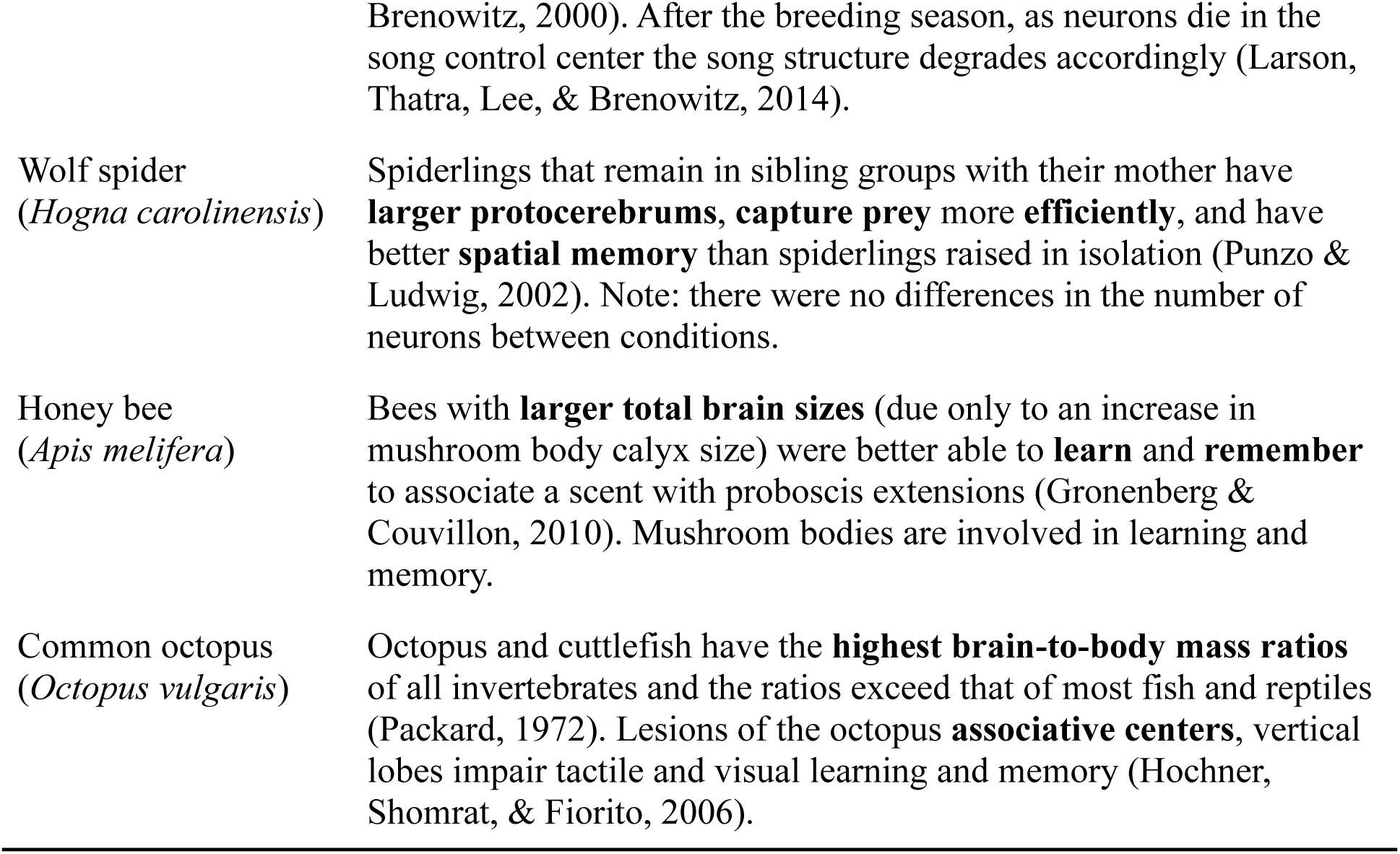
Examples of how behavior (directly tested) links with brain measures at the within-species level. These are the kinds of data that can contribute to the bottom-up approach to generate hypotheses based on validated data.

| Taxa | Description |
| --- | --- |
| Black-capped chickadee (*Poecile atricapillus*) | Birds in harsher environments (higher latitudes) had larger **hippocampus volumes** with more **neurons** (T. C. Roth & Pravosudov, 2009) and more **neurogenesis** (Chancellor, Roth, LaDage, & Pravosudov, 2011), were more efficient at **recovering caches** (spatial memory) and better at an **associative learning** task than conspecifics from less harsh environments (Pravosudov & Clayton, 2002). |
| Mountain chickadee (*Poecile gambeli*) | Individuals living at higher elevations had better **spatial memory** and more **hippocampal neurons**. Higher elevation environments are more challenging because variables such as day length and temperature vary more annually than they do at lower elevations (Freas, LaDage, Roth II, & Pravosudov, 2012). |
| Gambel's white-crowned sparrow (*Zonotrichia leucophrys gambelii*) | **Neurogenesis** increases in the **song control nucleus HVC** just prior to the breeding season (Larson et al., 2013). The breeding season is correlated with a higher **song quality** than in the non-breeding season (Meitzen, Thompson, Choi, Perkel, & Brenowitz, 2009; Tramontin & |

For example, spatial navigation behavior has been directly linked to the hippocampus using the bottom-up approach. Supporting evidence comes from intraspecies behavioral studies in birds with hippocampal lesions, which indicates the causal relationship between location memory and the hippocampus (Hampton & Shettleworth, 1996; Patel, Clayton, & Krebs, 1997). Additionally, ecological correlates were found in black-capped chickadees where individuals living in harsher environments (higher latitudes) were more efficient at recovering caches (spatial memory) and had larger hippocampal volumes with higher neuron densities and more neurogenesis than individuals at lower latitudes (Chancellor et al., 2011; Pravosudov & Clayton, 2002; T. C. Roth & Pravosudov, 2009). Further, real-time brain activity has been paired with navigational behavior in rats; when navigating through a maze, particular neurons (place cells) fired at particular locations in the hippocampus (Gupta, van der Meer, Touretzky, & Redish, 2010). Later, when the rats were not in the maze, rats mentally ‘ran’ through the maze and even invented novel routes as evidenced by the sequences of the firing of their place cells (Gupta et al., 2010). Place cell research and experimental designs that behaviorally test episodic memory (e.g., Clayton & Dickinson, 1998) provide evidence for brain-behavior causations from the bottom-up.

Where functional assays are either unfeasible or unethical, causality can be determined using a quantitative genetics approach to model how multiple measured traits are related to one another. Analyzing brain and behavioral data in pedigrees or full-sibling/half-sibling families allows the estimation of genetic correlations between traits (i.e., demonstrating variation in two traits that share a common genetic basis). If variation in brain size or composition causatively produces variation in behavior we should expect strong genetic correlations between these traits. This approach can be used not only to test brain-behavior relationships (e.g., Kotrschal et al., 2014), but also to help resolve debates about, for example, the relative roles of domain general and domain specific cognition (e.g., Pedersen, Plomin, Nesselroade, & McClearn, 1992), and developmental models of brain evolution (e.g., Hager, Lu, Rosen, & Williams, 2012; Noreikiene et al., 2015).

### 5.3 The comparative approach as a tool for generating hypotheses and testing generality

Although we argue for increased emphasis on intraspecific studies to validate causative relationships, the comparative approach will remain an integral part of investigations of the evolution of brains and cognitive abilities, though their scope and design might change. Phylogenetic studies extend and inform detailed intraspecific studies, ideally leading to constant feedback that can enhance both (Figure 3). Continuously developing comparative approaches have the potential to reduce noise from small sample sizes, reveal relationships among multiple interfering traits, and indicate the directionality of a causal association—though not all at once. Combining findings from multiple populations can inform mechanistic studies by illustrating the range of possible solutions that might exist, indicating where natural experiments might have shaped evolution in similar ways, revealing potential mediators by indicating in which taxonomic groups established relationships break down, and showing which species to target for further study. In particular, the systematic combination of effect sizes from population studies in phylogenetic meta-analyses reduces noise and can test the robustness of an association between brain measures and behavior while also revealing potential mediators that systematically change the form of the association in some populations or species (Nakagawa & Santos, 2012). For example, they might reveal whether the heritability of brain measures might depend on environmental variability.

**Figure 3:**
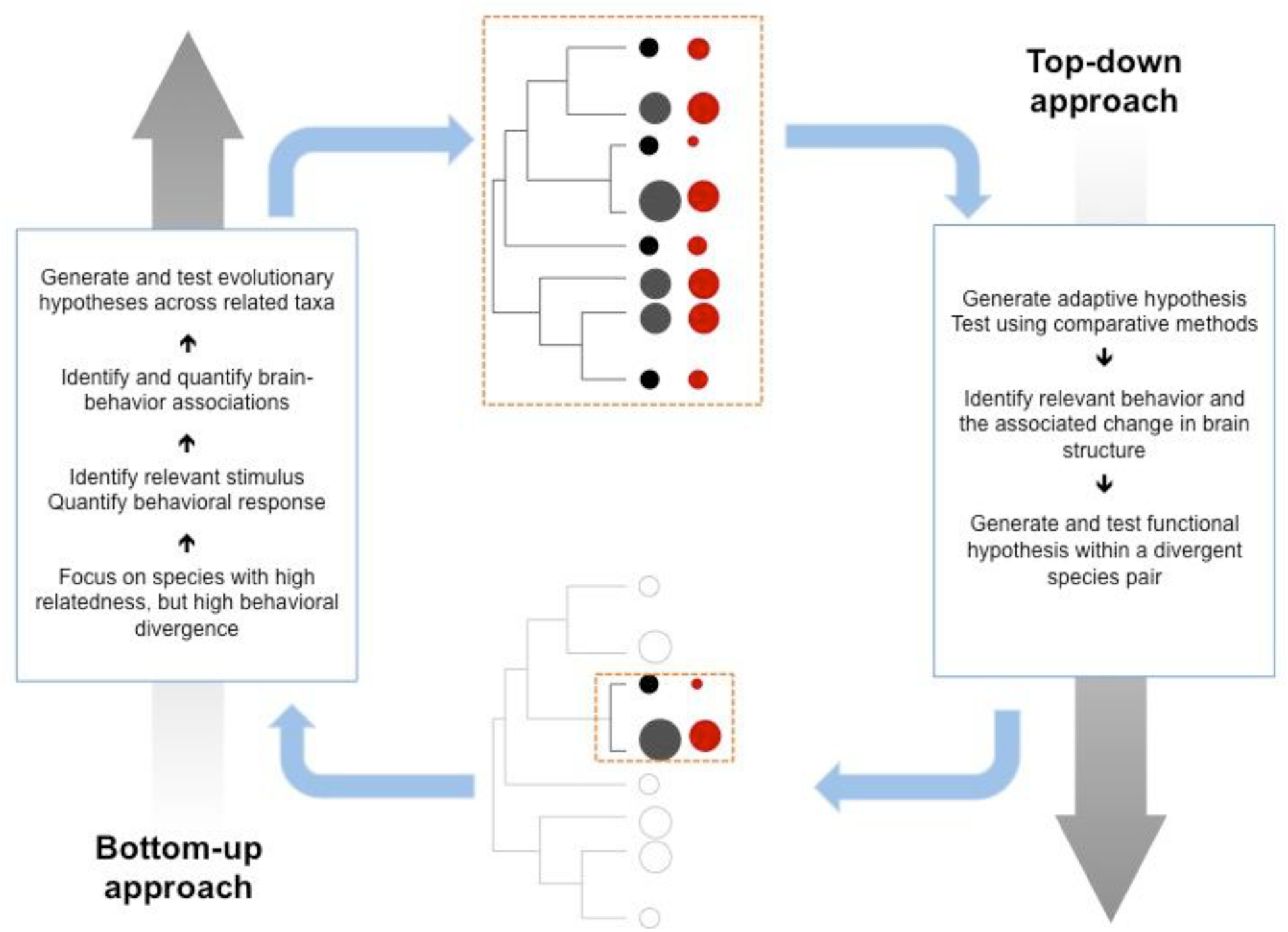
Integrating the top-down and bottom-up approaches.

In turn, the historical component of phylogenetic reconstructions extends population-level studies by revealing whether detected patterns are evolutionarily stable or lineage-specific, and they can contribute to determining causal or adaptive relationships between traits by revealing temporal contingencies (Beaulieu, Jhwueng, Boettiger, & O’Meara, 2012; Pagel, 1999; Pagel & Meade, 2006) in whether a behavior consistently changed prior to or after associated changes in brain measures. The historical component provided by phylogenetic comparisons is necessary to determine whether traits not only occur together, but whether they evolved together. For example, while the enlarged brains (compared to most other reptiles) among birds appear to be linked to cognitive capacities required for flight (Balanoff, Bever, Rowe, & Norell, 2013), evolutionary origins of flying behavior are not associated with particular increases in endocranial volume (Balanoff, Smaers, & Turner, 2016).

Our discussion of the power of the comparative approach in elucidating the adaptive history of traits indicates the inherent limits in fully explaining traits that supposedly make any species unique. The evolutionary processes themselves are not unique, but the particular combination of processes at play are. As such, understanding how such processes come together in a particular instance is problematic due to a lack of evidence required to confirm these hypotheses (Tucker, 1998). In addition, studies that focus on extraordinary traits in a single species (such as humans) frequently risk misrepresenting evolutionary processes by fixating on the endpoint as an optimal solution, whereas evolution typically progresses by responding to stochastic variation in selection regimes, incrementally adapting to the environment.

### 5.4 Scaling across taxa to integrate evidence

The bottom-up approach we suggest means that scaling across taxa will initially be more difficult to achieve because studies will have to be designed to take into account the characteristics of the particular species, as well as its phylogenetic and ecological context. Questions, approaches, and methods might need time to converge or to be repeated across a relevant sample of different taxa (Figure 3). However, over an intermediate time frame, the bottom-up approach will be invaluable for comparing and elucidating brain and cognitive evolution across taxa. Although the bottom-up approach initially makes scaling look difficult, we think it has two advantages. First, rather than positing or assuming a coarse-grained, cross-taxa category and applying it across a range of cases (thus losing ecological relevance and increasing the potential for *post-hoc* explanations and reification), the bottom-up approach makes scaling a much more piecemeal, empirically tractable matter. Second, it more easily allows scaling to take place in an evolutionary context. Understanding whether the same genes, genetic pathways, neural regions, neural physiology, and/or neural networks are involved in generating cognitive abilities across taxa will provide us with an understanding of how evolution has shaped the diversity of brains and the behavior they produce. In this sense, phenotypic heterogeneity and taxonomic diversity become a tool for discovery, rather than a source of statistical noise.

It is not straightforward to bring together the disparate evidence involved in shifting from local experimental contexts to cross-taxa, evolutionary hypotheses. However, a detailed understanding of the mechanisms underlying brain measures and behavior is crucial to clarify whether traits are homologous, analogous, or completely independent solutions to ecological challenges. To give a sense of the possibilities for integration, we sketch three kinds of approaches to shifting from local (bottom-up) to general (top-down) scales (from Currie, 2013; see also Mikhalevich et al., 2017):

1. Detecting homologous relationships, where the same brain measures and behavior are related to the same environment across species descended from a recent common ancestor (Currie, 2012), offers opportunities to combine independent findings into one mechanistic pathway. In these instances, inheritance and stabilizing selection have maintained a stable trait, such that findings from one species can be accurately inferred for another. Such investigations will rely on integrated models that bring together disparate evidence to support hypotheses about the evolutionary, developmental, and ecological features of a particular lineage.
2. Determining whether the same behavior occurs in similar environments across distantly related species can indicate environments most likely relevant for the emergence of the behavior. A bottom-up approach can reveal whether the observed behavior represents analogous re-emergence of a behavior within the same adaptative environment (e.g., repeated evolution of feathers across dinosaurs; B. K. Hall, 2003; McGhee, 2011). This approach will rely on parallel models that identify brain-behavior correlations within related taxa for which the main principles of brain evolution are known to be similar. As closely related taxa will likely share meaningful brain-behavior correlations, such models are likely to be well-validated, stable, and causally meaningful.
3. Observing a similar behavior in similar adaptive environments can reveal whether the behavior represents a convergent solution to the same environment (e.g., feathered wings for flight versus bat wings) or whether the relationship is more complex (e.g., wings to escape into the air versus jumping legs; Currie, 2014; Pearce, 2012; Powell, 2007). This type of convergent model is similar to the top-down approach; however, convergent modeling avoids many of the cross-taxa comparison problems by i) being placed in an explicitly ecological and phylogenetic context, ii) being carried out alongside parallel and integrated modeling, and iii) avoiding over-interpretation that arises from defining categories based on superficial similarities because convergent models are inherently sensitive to the explanatory limits of analogous categories (see Griffiths, 1994 for a discussion of these limits).

There are a wide variety of different scales at which we may need to infer evolutionary relationships between brain measures, behavior, and environments across taxa, and the ecological and evolutionary relevance granted by starting in local contexts is crucial for doing this.

## Conclusion

We support a two-pronged strategy for understanding cross-taxa relationships between brain size, brain composition, behavior, and cognition that focuses on ecologically relevant contexts rather than attempting broad scale comparisons at gross phenotypic levels. The first prong is an experimental program examining correlations in closely related species; the second prong involves the piecemeal identification of correlations at broader taxonomic scales. We have contrasted our approach with one that has become dominant in recent years. The alternative approach relies on coarse-grained phenotypes and proxy-measures, typically in anthropocentric contexts, and attempts to apply these to cross-taxa, correlative contexts. We have highlighted a number of limitations to this approach. First, applying anthropocentric conceptions of brain correlates with behavior to disparate taxa comes at the crucial cost of ecological and evolutionary coherence. Second, the heterogeneity of brain composition and behavior makes coarse-grained conceptions problematic because cross-taxa comparisons inevitably discount variation that matters for particular lineages. This variation creates noise in statistical comparisons. Heterogeneity can also be a source of interference because various interdependencies both between brain structures (e.g., in development or function) and between multiple behavioral and ecological traits undermine our capacity to identify selection pressures shaping individual traits or systems. Third, beginning with ‘general’ measures of intelligence potentially leads to reification and the establishment of misguided or causally meaningless properties. The top-down approach has not necessarily been misguided itself: scientific progress is often facilitated by applying relatively crude measures, highlighting the value of using many investigative techniques. Indeed, the heterogeneity of these traits have become known to us *because* the top-down approach has exposed inconsistencies through cross-species correlations. However, it is time to take the cognitive, behavioral, and brain features of particular lineages seriously, rather than demand that they be shoehorned into anthropocentric notions, or judged against some general metric. In doing so, a more general understanding of the nature of cognition and behavior, and their relationship with brain measures will be built from the bottom up.

## Acknowledgements

We are grateful for manuscript feedback from Nicky Clayton, Tim Clutton-Brock, and Rob Barton, and we appreciate Christian Rutz for discussions about New Caledonian crow foraging innovations. We thank our funders: the Isaac Newton Trust and Leverhulme Trust for a Leverhulme Early Career Fellowship to CJL, which funded the workshop on which this paper is based; NERC for an Independent Research Fellowship to SHM; the European Research Council (Grant No. 3399933; SAJ); the Royal Society for a Dorothy Hodgkin Research Fellowship to NJB; the Royal Society of New Zealand Marsden Fund (UOC1301; FRC); the National Science Foundation (NSF BCS 1440755; RM); the John Templeton Foundation (AB); and the Templeton World Charity Foundation (AC; note: the opinions expressed in this publication are those of the author(s) and do not necessarily reflect the views of Templeton World Charity Foundation).

## Authorship Contributions

All authors contributed to the concepts in this paper at a workshop organized by Logan in March 2017. Montgomery structured the paper. All authors wrote and edited the paper. Logan, Montgomery, Boogert, and Mares served as managing editors. All authors approved the final version for submission. We certify that what we have written represents original scholarship.

## Box 1. What can comparative cognitive tests tell us?

Performance on cognitive tests arguably offers the most direct measure of ‘*intelligence*’ and can therefore be seen as an improvement over other behavioral measures – such as innovation rate – which have been used as proxies for intelligence. However, cognitive abilities can still only be *inferred* from test performance, rather than directly measured. Performance on any given test will depend on a suite of abilities and behaviors (such as motor abilities, perception, attention, motivation, fear), beyond the cognitive ability in question, which mean that successes or failures can occur for a range of different reasons across subjects (Rowe & Healy, 2014; A. Seed, Seddon, Greene, & Call, 2012).

Let’s take the example of MacLean and colleagues’ (2014) collaborative study, comparing brain size with measures of self-control across species. Here, 36 species of mammals and birds received two cognitive tasks: the A-not-B task and the cylinder task. In the A-not-B task a human demonstrator places food in cup A multiple times. Once the subject has retrieved a reward from cup A three times in a row, they are given a test trial where food is first placed in cup A, and then visibly moved to cup B. Subjects have to inhibit choosing previously rewarded cup A to succeed. This is a commonly used developmental task, on which babies under 10 months typically display perseverative reaching errors (Smith, Thelen, Titzer, & McLin, 1999). However, in addition to ‘self-control’, to pass this task, subjects also need to be capable of accurately tracking the movement of food by human hands. This is trivial for humans, and may also be relatively easy for great apes or other primates, given that they are closely related to humans and also possess hands. However, there is growing evidence that many species struggle to use this type of information from a human demonstrator (e.g., Erdőhegyi, Topál, Virányi, & Miklósi, 2007; Shaw, Plotnik, & Clayton, 2013); thus, this task may have systematically disadvantaged certain species for reasons unrelated to their self-control (Jelbert et al., 2016). Crucially, in MacLean and colleagues’ study, although some animals (lemurs, dogs and pigeons) were explicitly trained to select experimenter-baited containers during pre-training, most other species were not. All non-primates that were given both tests, and had *not* been trained to attend to the demonstrator, performed substantially worse on the A-not-B task than on the cylinder task. Elephants, given the A-not-B task only, failed every trial. Without knowing whether subjects possessed one of the key requirements for the A-not-B test, we cannot know whether their poor test scores actually reflect poor self-control. In a study that directly tested for this effect, New Caledonian crows that had been explicitly trained to attend to a human demonstrator went on to pass 67% of A-not-B test trials, while a control group, trained on an unrelated inhibitory control task, passed only 7% (Jelbert et al., 2016).

The particular use of the A-not-B task is a clear example of a situation in which comparing cognitive test scores from different species, and relating them to brain size, is unlikely to provide us with a meaningful comparison of the cognitive ability in question. Given the range of factors that can influence test success, the majority of cognitive tests will suffer from limitations like this to some degree. For example, in MacLean’s cylinder task, one key variable that would influence performance was the amount of experience different subjects had with transparent materials. In the cylinder task, subjects first learn to retrieve a reward from the open end of an opaque cylinder, and are then presented with a transparent cylinder, where the reward is now visible through the tube. Subjects are considered to have failed the task if they touch or peck the front of the transparent tube, rather than detouring to the open side. Kabadayi et al. (2016) highlighted that a number of bird species showed learning effects over the course of 10 trials in the cylinder task, suggesting that relative unfamiliarity with transparent Perspex might have contributed towards this behavior.

Other factors that might have influenced performance can also be hypothesized. These could include morphological differences (i.e., performance in the cylinder task may vary depending on whether a subject must insert their arm or their head into the tube to obtain the reward), perceptual differences (i.e., how well can different subjects perceive the transparent material?), behavioral differences (i.e., does this species typically explore or avoid new objects?) or motivational differences (i.e., to what extent is the animal motivated to directly obtain the reward?) or any other task specific variables. Thus, while here we have highlighted some specific examples, performance on any cognitive test will be influenced by numerous sources of variation, both across individuals and across species, in addition to the cognitive ability in question.

To address this, minimizing any unnecessary task demands (such as the use of human demonstrators) is the first step in designing comparative tests. It is also crucial to level the playing field from the bottom up, including baseline criteria training to ensure that all subjects meet specific key requirements, before the test of the desired cognitive ability begins. Focusing on groups of more closely related species will also help to limit the number of ways in which subjects’ performance could vary beyond the ability in question.

## Box 2. Spider behavior varies according to environmental differences

Traditionally, vertebrates have been used as subjects to help answer probing questions relating to animal cognition, including studies of working memory, search images, expectancy violation and insight (e.g., Kamil & Bond, 2006; Köhler, 1924, p. 192; Pepperberg & Kozak, 1986; Shettleworth, 2010). However, recent research demonstrates that invertebrates, despite having much smaller brains, perform similarly on cognitive tasks (Jakob, Skow, & Long, 2011; Perry, Barron, & Chittka, 2017), thereby challenging notions of brain size translating into cognitive ability.

Jumping spiders (family Salticidae), for example, have been of considerable interest in the context of cognition, partly because they have large forward-facing principal eyes that play a major role in high-precision visual discrimination (Harland, Li, & Jackson, 2012), supporting tasks such as selective attention and planning (Jackson & Cross, 2011). There is still much to be learned about spider neurobiology (but see Menda, Shamble, Nitzany, Golden, & Hoy, 2014); however, excellent vision may be part of the solution for how an animal with a minute brain can perform cognitive tasks in its environment (e.g., Pfeifer & Iida, 2005).

Moreover, spiders are an excellent system for investigating how variation in cognitive abilities may be associated with ecology. The salticid genus *Portia* has provided us with many insights because the species from this genus eat other spiders (Jackson & Wilcox, 1998) and deploy a variety of strategies to avoid being eaten by their prey (Jackson & Cross, 2011). For example, when at the edge of another spider’s web, *Portia* is known to deploy a specialized strategy of moving its eight legs and two pedipalps across the silk in ways that may mimic the movements made by a trapped insect (Jackson & Cross, 2013). These web signals are generated using trial and error (Jackson & Wilcox, 1993); *Portia* repeats a signal when the resident spider starts moving toward *Portia* across the web, and changes to using a different signal when it does not succeed at eliciting an approach by the resident spider.

However, differences in the use of this tactic have been observed in different populations of the same species. In the Philippines, two populations (Los Baños and Sagada) of *Portia orientalis* (formerly *P. labiata*) encounter different spider species as prey. The site at Los Baños is a low-elevation tropical rainforest where *P. orientalis* encounters a wider variety of prey species, including species that are particularly dangerous, such as *Scytodes pallidus*, a spitting spider that specializes on salticids as prey (Li, Jackson, & Barrion, 1999). In contrast, the site at Sagada is a high-elevation pine-forest where *P. orientalis* encounters a smaller number of prey species, and does not encounter the particularly dangerous prey species that are present in Los Baños (Jackson & Carter, 2001; Jackson, Pollard, Li, & Fijn, 2002). For the Los Baños *P. orientalis*, a greater reliance on flexible predatory strategies, such as using trial and error when generating signals in other spiders’ webs, is likely to be of greater importance than for the Sagada population. Indeed, when at the edge of another spider’s web, the *P. orientalis* from Los Baños repeated a signal that elicited movement by the resident spider significantly more often than the Sagada population, and were also more likely to change a signal when it did not elicit movement by the resident spider (Jackson & Carter, 2001).

Individuals from the Los Baños population also learned faster when faced with a novel situation of escaping from an island in a water-filled tray (Jackson, Cross, & Carter, 2006). After leaving the island, the spider first needed to reach an atoll before it could then reach the edge of the tray, but the distance was too far for the spider to clear by leaping alone. Instead, the spider could first reach the atoll by swimming across or, alternatively, it could leap before swimming the rest of the way. However, before a trial began, the researchers decided at random which of these two tactics (leaping or swimming) was the ‘successful’ tactic for reaching the atoll. If the spider used the successful tactic (e.g., when the successful tactic was leaping and the spider leaped), the researchers used a plastic scoop to make small waves in the water to help the spider across to the atoll. The spider then made its next move from the atoll. If, however, the spider used the ‘unsuccessful’ tactic (e.g., when the successful tactic was swimming and the spider leapt), the researchers used the plastic scoop to force it back to the island. To successfully reach the atoll, the spider then needed to switch to using the other tactic on its next attempt from the island. Similar to when making signals in webs, individuals of the Los Baños population repeated tactics when successful, and switched tactics when unsuccessful, significantly more often than the Sagada individuals (Jackson et al., 2006).

It is currently unknown whether such observed differences in arachnid behavior are causally related to neural architecture. However, applying the bottom-up approach to the study of cognition in this taxon will likely contribute substantially to our understanding of how cognition relates to ecology and neurobiology. As well as showing a wide variety of foraging strategies in various environments, spiders are relatively easy to study in the field and laboratory, especially when compared with large vertebrates.

